# A wheat immune receptor pair executes cell death through a helper MLKL

**DOI:** 10.64898/2026.08.07.743466

**Authors:** Josh W Bennett, Yu Sugihara, John F Haidoulis, Clémence A Rodney, Rafał Zdrzałek, Enzo Zanchet, Indira Saado, Pirita Paajanen, Paul Nicholson, Soichiro Asuke, Mark J Banfield

**Affiliations:** Department of Biochemistry and Metabolism, John Innes Centre, Norwich Research Park, Norwich, NR4 7UH, UK; Department of Genomics and Breeding, Iwate Biotechnology Research Centre, Kitakami 024-0003, Japan; Department of Computational and Systems Biology, John Innes Centre, Norwich Research Park, Norwich, NR4 7UH, UK; Department of Crop Genetics, John Innes Centre, Norwich Research Park, Norwich, NR4 7UH, UK; Graduate School of Agricultural Science, Kobe University, Kobe 657-8501, Japan

## Abstract

To promote disease resistance, plant nucleotide-binding, leucine-rich repeat (NLR) immune receptors often require paired co-receptors. In many cases, paired NLRs comprise one NLR to perceive effectors (the sensor) and another NLR to execute cell death (the helper). However, NLRs can also pair with sensor kinase fusion protein (KFP) receptors, but whether non-NLR components within such pairs can execute cell death, remains unclear. Here, we investigate the mechanism of an immune receptor pair comprising the wheat NLR <u>R</u>wt<u>3</u>.6.8 <u>NLR</u> (R3NLR) and an MLKL protein, <u>R</u>wt<u>3</u>.6.8 <u>a</u>ssociated <u>k</u>inase (R3AK). Using *Nicotiana benthamiana* transient expression assays we confirmed that both R3NLR and R3AK are required for cell death in response to blast pathogen effectors PWT3, PWT6 or PWT8. Through mutational analysis we show the 4-helical bundle (4HB) domain of R3AK is required to execute cell death and R3AK can be made auto-active by perturbing the kinase catalytic active site. Activation of R3AK is also associated with a shift to a higher oligomeric state. Furthermore, as the NLR R3NLR is not actively involved in the execution of cell death we hypothesise that R3AK acts as a helper. A phylogenetic analysis indicates widespread distribution of this paired configuration in Poales. Together, this study establishes a novel resistance mechanism involving a non-canonical NLR/MLKL system.

**Significance Statement:** Here we investigate the mechanism of a novel plant immune receptor pair from wheat, R3NLR/R3AK. A nucleotide-binding, leucine-rich repeat (NLR) receptor and a mixed lineage kinase like (MLKL) protein are both required to mediate resistance to blast pathogen effector proteins PWT3, PWT6 and PWT8. Adopting a mutagenesis approach, we show that the MLKL protein executes cell death through its N-terminal 4-helical bundle domain, and this is associated with a shift to a higher oligomeric state. Mutations of conserved sequence motifs in the NLR support its role as a sensor, although effector interactions have not yet been observed. This study reveals a plant immune receptor pair that functions via a putative NLR sensor paired to a cell death executing MLKL protein.

## Introduction

Crop yield losses to plant disease are a major threat to agricultural productivity (1). The causative agent of rice and wheat blast, *Magnaporthe oryzae* (syn. *Pyricularia oryzae*), is particularly devastating, leading to disease on a range of cereal crops. Infection is facilitated by a repertoire of secreted virulence proteins called effectors (2–6). These proteins can be recognised by plant immune receptors and understanding the mechanism of receptor function is important for enabling effective resistance to improve global food security (7). One class of well-studied intracellular immune receptors are the nucleotide-binding leucine-rich repeat (NLR) proteins (8). Plant NLRs typically comprise three domains: an N-terminal executor domain, an NB-ARC activation/oligomerisation domain and a C-terminal leucine-rich repeat (LRR) (9). NLRs are often classified into three groups depending on their N-terminal executor domains; comprising either a 4-helical bundle (4HB) coiled-coil (CC) domain (CNLs), a resistance to powdery mildew-8 (RPW8) domain, or a Toll/Interleukin-1 (TIR) domain (TNLs), although the latter group is absent in monocot species. Activation of N-terminal executor domains typically leads to programmed cell death, preventing pathogen spread (10). Effector perception by NLRs can either be direct, where the effector binds to the LRR or an integrated decoy domain, or indirect, where the NLR perceives modifications or binding of effectors to a separate host target (11–16).

Some NLRs function as autonomous units mediating both effector perception and initiation of immune responses; they are often referred to as singleton NLRs (17). However, NLRs also work in genetically linked, co-expressed pairs comprising a ‘sensor’ NLR to perceive an effector and a paired ‘helper’ NLR which executes cell death (18). Nevertheless, upon activation (effector perception) NLRs typically organise into higher order complexes (resistosomes) and their α1 helix can insert into cell membranes to form channels (CNLs/RPW8) or to produce small molecules for signalling (TNLs) (19, 20).

Alongside the well characterised NLRs, kinase fusion proteins (KFPs) are an emerging class of plant intracellular immune receptors, especially in Poaceae. The majority of KFPs reported to date are tandem kinase proteins (TKPs) (Sr60, Un8, WTK4, Rpg1, RWT4, Sr62, Yr15, Lr9, Pm57, Rmo2 and RWT7 (21–30). However, kinase proteins fused to other domains have also been reported, such as the 4HB (also referred to as a HeLo domain) domain (Pm13), C2 transmembrane domain (Pm4) or multiple DUF domain (Sr43) kinase fusions (31–34). TKPs mainly consist of two kinase/pseudokinase domains, but can also contain additional diverse protein domains, which may have roles in effector perception (27, 35–37). Recently, the TKPs Sr62^TK^ and RWT4 were shown to be genetically linked to an NLR which is required for downstream signalling (29, 38), suggesting a new model for paired immune receptors with ‘sensor’ KFPs and paired ‘helper’ NLRs. In this model, the C-terminal pseudokinase domain is repressed under normal conditions via intramolecular interactions which are released when the cognate effectors bind to the N-terminal kinase domain. The pseudokinase then activates the paired NLR through direct interaction to execute cell death (39, 40). Interestingly, other KFPs, such as Pm13, a mixed lineage kinase like (MLKL) protein, have been shown to function as direct executors of cell death (32) and may operate as singletons or through linkage with paired NLRs (41).

MLKLs comprise an N-terminal 4HB domain and a C-terminal kinase/pseudokinase domain, linked by a brace region (42). In mammals, activated MLKLs form oligomers and permeabilise the cell membrane via their 4HB domain, leading to cell death (43, 44). MLKLs are also present in plants and are highly conserved in angiosperms (45, 46). The *Arabidopsis thaliana* MLKL, *At*MLKL2, can be activated by TIR-signalling to form a calcium channel, executing cell death akin to CNLs (46, 47). By contrast, transient expression of the brace and kinase domain of the phylogenetically distinct MLKL protein Pm13 alone was sufficient to cause cell death in *N. benthamiana* (32). The discovery of Pm13 led to the identification of more MLKLs enriched in *Triticeae* arising from multiple independent fusion events (32, 33). This apparent convergent evolution demonstrates the importance of MLKL domain architecture in plant immunity. However, how other distinct plant MLKLs function is unknown.

*Rwt3* encodes a wheat NLR that recognises the *M. oryzae* effector PWT3, a genetic interaction that underpinned a host jump leading to the wheat blast pandemic (3, 48). Recently, the *Rwt3* resistance locus has been shown to comprise not only an NLR (24), but a genetically paired NLR and KFP in a head-to-head orientation (49). Furthermore, this immune receptor pair has previously been identified as the resistance locus recognising the avirulence effector PWT6, formerly dubbed *Rwt6* (49, 50). Intriguingly, both genes are required for resistance to the *Magnaporthe oryzae* (*M. oryzae*) effectors PWT3 and PWT6 and also a third effector, PWT8 (49). Therefore, this resistance locus is now termed *Rwt3.6.8* and encompasses both a CNL named <u>R</u>wt<u>3</u>.6.8 <u>NLR</u> (R3NLR) and an MLKL protein, <u>R</u>wt<u>3</u>.6.8 <u>a</u>ssociated <u>k</u>inase (R3AK), establishing a novel NLR pairing involving a KFP. However, the mechanistic basis of how these immune receptors contribute to resistance is still unknown.

In this study, we built on the discovery of R3NLR/R3AK resistance to investigate the molecular mechanism of how this NLR/MLKL pair executes an immune response to three sequence and structurally diverse *Magnaporthe* effectors (49). We demonstrate that the R3NLR/R3AK system can be recapitulated through transient expression in *N. benthamiana* and both R3NLR and R3AK are required for cell death when co-expressed with effectors. Using a mutagenesis approach, we explored the roles of well-established sequence motifs in R3NLR and R3AK for underpinning cell death activity. We demonstrate that the 4HB domain of R3AK, but not the R3NLR CC domain, is required for cell death on co-expression with effectors. Furthermore, we obtained an auto-active variant of R3AK that is sufficient to cause cell death in the absence of R3NLR or any effectors. We also identify that an ‘active’ R3AK mutant forms higher order oligomers than in the resting (‘inactive’) state. Based on this data, we suggest that in this paired system the MLKL, R3AK, may act as the ‘helper’ executing immune-related cell death. Using a phylogenetic approach, we also show that genetic NLR/MLKL pairings are present in Poaceae and Zingiberales, suggesting they are not limited to *Triticum aestivum*.

## Results

### R3NLR/R3AK mediates cell death when co-expressed with three structurally diverse *Magnaporthe* effectors in *N. benthamiana*

R3NLR (NLR) and R3AK (MLKL), form a genetically linked head-to-head pair (**Fig. 1a**) required for resistance against three *M. oryzae* effectors PWT3, PWT6 and PWT8 (24, 49, 50). Since NLR mediated resistance often involves cell death, we tested whether this could be recapitulated in *N. benthamiana* by transiently co-expressing R3NLR and/or R3AK with cognate effectors PWT3, PWT6, PWT8 or the non-recognised effector AVR-PikD as a negative control. All proteins accumulated in plant tissue as determined by western blotting (**Fig. S1**). Co-expression of R3NLR and R3AK without effectors, or with the non-cognate effector AVR-PikD, resulted in low levels of cell death indicating some auto-activity when transiently expressed in *N. benthamiana* (**Fig. 1b-d**). Importantly, co-expression of either R3NLR or R3AK alone with each effector was not sufficient to induce cell death whereas co-expression of R3NLR and R3AK with either PWT3, PWT6 or PWT8 gave significantly stronger levels of response (**Fig. 1b-d, S2**). These data are consistent with previous findings showing that both R3NLR and R3AK are necessary for immune signalling in response to recognition of PWT3, PWT6 and PWT8 (49).

**Figure 1:**
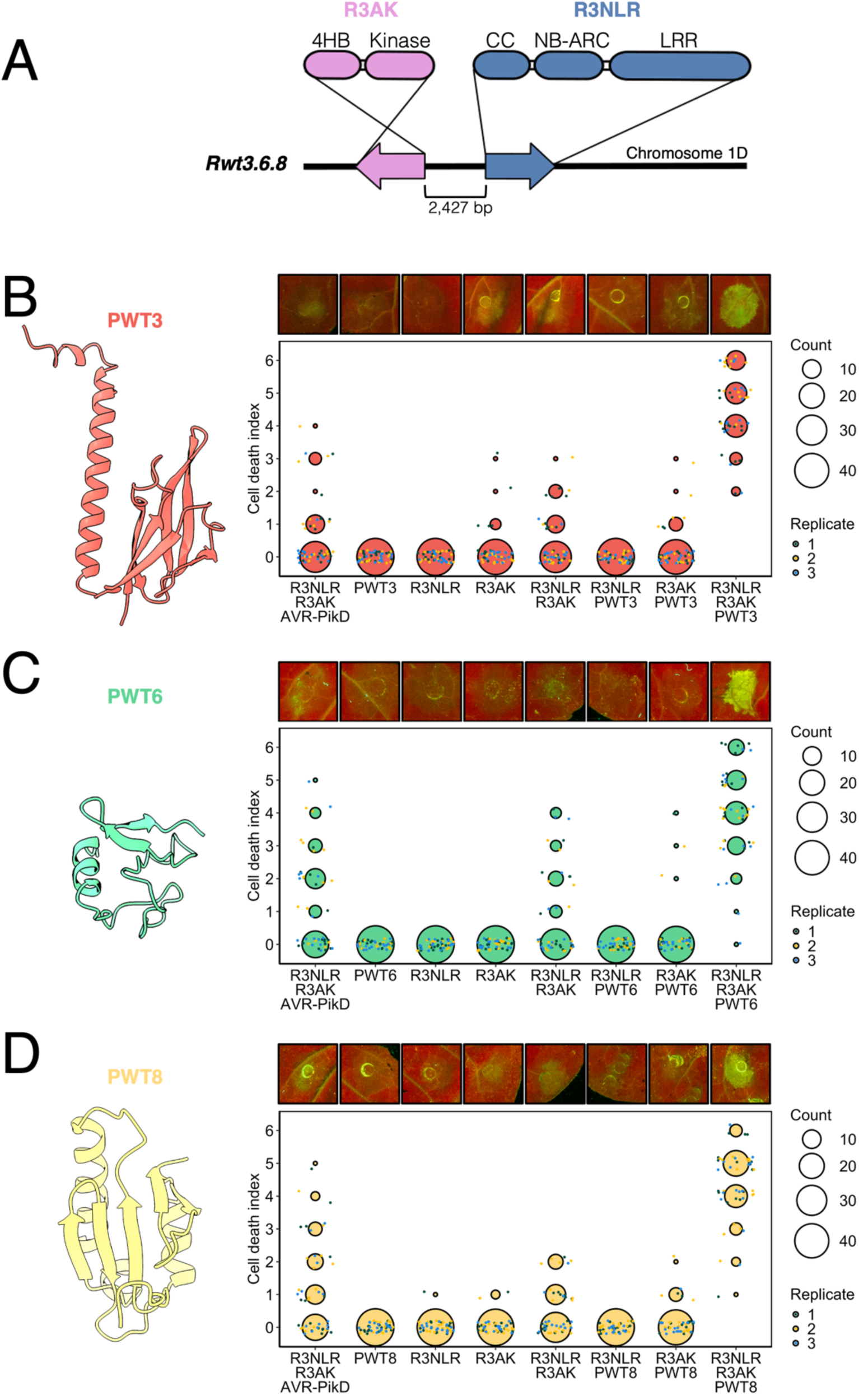
A genetically linked wheat CNL and MLKL are necessary to respond to three sequence and structurally diverse *M. oryzae* effectors in *N. benthamiana*. (A) Schematic diagram of the head-to-head pairing of R3NLR and R3AK and their respective domains architectures. (B-D) Effector structures (predicted by AF3, PWT3 (B), PWT6 (C), or crystal structure, (PWT8 (D)) are shown alongside representative abaxial side leaf images taken under UV light (top) and dot plots showing scoring for the cell death across repeats. For the cell death plots, each dot represents an individual scoring point, which are randomly jittered for ease of visualisation (dot colour represents the different biological replicates). The size of the larger circles represents the number of times this score was observed, besthr statistical analysis is shown in Fig. S2.

PWT3, PWT6 and PWT8 share low amino acid sequence homology **(Fig. S3, Table. S1)**, which prompted investigation of their structures. We were unable to obtain crystal structures of PWT3 and PWT6 and used AF3 to predict their conformations. The fold of PWT3 is confidently predicted (**Fig. 1b, S4**) and TM-align (51) revealed significant structural similarity to the MAX fold effector AVR-Pia (PDB: 9RSV) (52), but with a C-terminal alpha helical extension (**Fig. 1b, Fig. S4, S5**). By contrast, the AF3 prediction of PWT6 is mostly disordered and of low confidence (**Fig. 1c, S4**). For PWT8, we were successful in solving the crystal structure using a selenourea based single-wavelength anomalous diffraction (SAD) approach (53), followed by refinement to 1.3 Å resolution (PDB: 31KJ). The structure revealed PWT8 adopts a β1-α1-α2-β2-β3-α3-β4-β5 topology, unlike previously identified effector folds (**Fig. 1d, Table. S2, S3**). These structural analyses suggest that R3NLR/R3AK can perceive sequence diverse effectors of different structural topologies.

We hypothesised that the R3NLR NLR may act as a guardee/sensor of the effectors binding to R3AK (kinase domain), in a mechanism similar to RWT4 (30). To explore this, we tested for interactions between all cognate effectors and both immune receptors using a co-IP (co-immunoprecipitation) approach. This included expressing R3NLR or R3AK alone or co-expressing both R3NLR and R3AK with effectors **(Fig. S6, S7).** While the well-established interaction between the NLR Pikp-1 and its effector AVR-PikD was observed (54), we did not detect binding of either R3NLR or R3AK to PWT3, PWT6 or PWT8. Therefore, whether R3NLR and R3AK are sufficient to directly bind these three effectors remains inconclusive.

### Effector-dependent cell death requires the R3NLR LRR domain but not the CC domain

We explored the molecular mechanism of the R3NLR/R3AK immune receptors through mutagenesis. For these experiments we focused on PWT3 as we had previously observed that it was the most robustly expressed in *N. benthamiana*. Firstly, we explored the roles of well-established sequence motifs in R3NLR for promoting cell death. An R3NLR mutant, R3NLR^3E^ (in which Leucine 12, Valine 15 and Leucine 19 are mutated to Glutamates), was generated that disrupts the N-terminal α1 helix of the CC domain, which typically abolishes cell death activity without compromising receptor activation (55). Surprisingly, when we sought to activate the R3NLR^3E^/R3AK system by co-expressing with PWT3, effector triggered cell death was still robustly observed (**Fig. 2a, S1 and S8a**). This result was also seen with a complete CC domain truncation of R3NLR (**Fig. S9**). These results indicate that execution of cell death in the R3NLR/R3AK system is not dependent on the CC domain of R3NLR.

**Figure 2:**
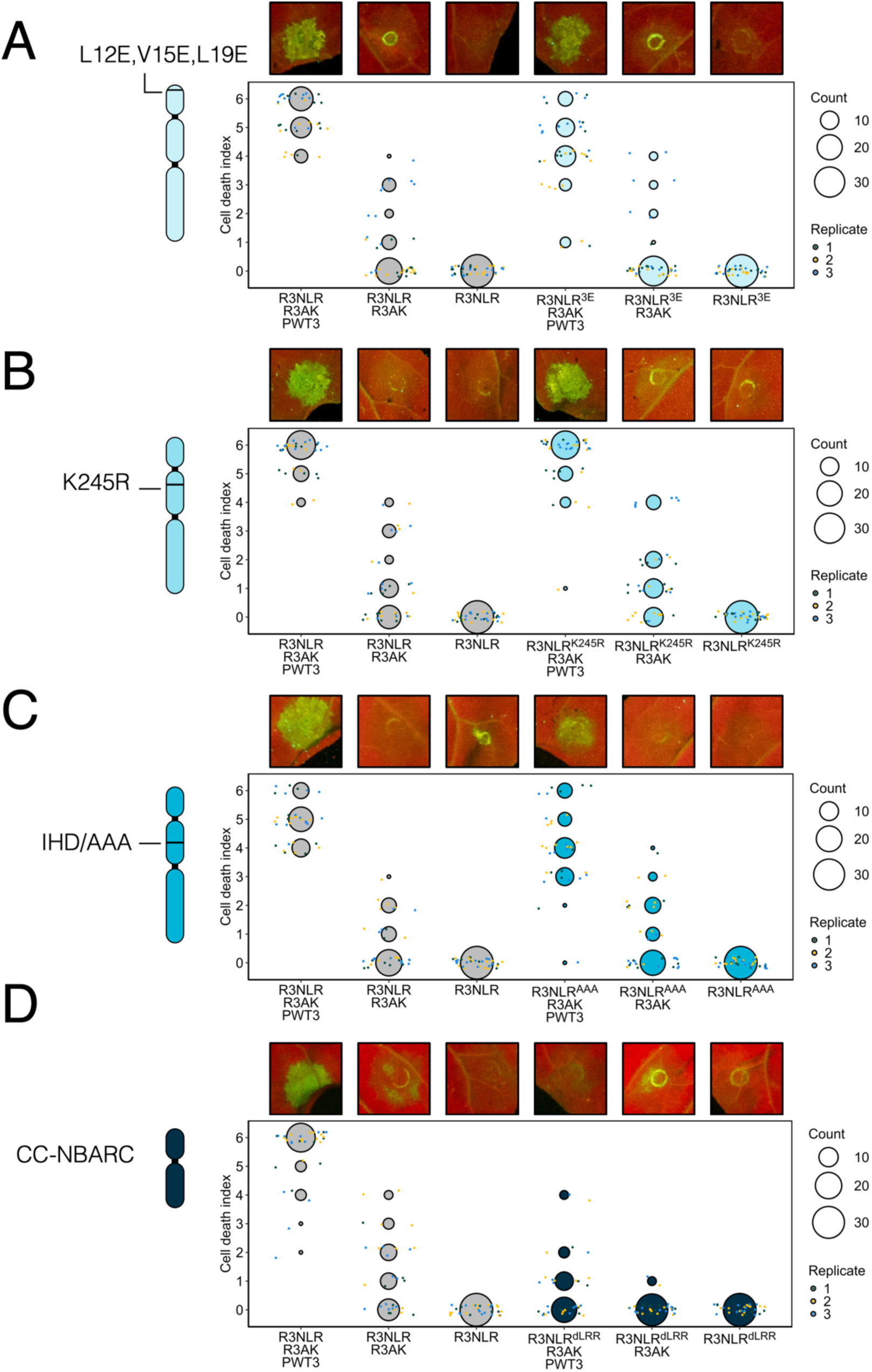
Cell death assays with mutants of R3NLR suggests this NLR does not behave as a helper NLR. Schematic diagrams highlighting the location of mutants within the R3NLR domain structure (left) are shown alongside representative abaxial side leaf images taken under UV light (top) and dot plots showing scoring for the cell death across repeats (bottom). (A) N-terminal CC domain mutants within the α1 helix, (B) “P-loop” mutant R3NLR^K245R^ within the NB-ARC domain, (C) “MHD” mutant R3NLR^AAA^ within the NB-ARC domain and (D) LRR domain truncation R3NLR^dLRR^. Details of the dot plots are as for Fig. 1, with the besthr statistical analysis shown in Fig. S8.

Mutations in the conserved P-loop and ‘MHD’ motifs of the NB-ARC domain typically render NLRs constitutively inactive or active (13, 56, 58, 59). To investigate the role of the R3NLR NB-ARC domain in the activation of the R3NLR/R3AK system we introduced mutations into the conserved P-loop and ‘MHD’ (IHD in R3NLR) motifs. For the P-loop mutant we generated a R3NLR^K245R^ construct. An ‘MHD’ mutant was generated by mutating the motif to triple Alanine (I527A, H538A, and D539A) forming a R3NLR^AAA^ construct. When expressed alone or co-expressed with R3AK and/or PWT3 in *N. benthamiana*, we did not observe any difference in the cell death response compared to wildtype R3NLR (**Fig. 2b, 2c, S1, S8b and S8c**). This indicates that R3NLR does not require intact P-loop or MHD motifs for cell death activation, in contrast to other previously characterised singleton or helper NLRs.

Collectively, as mutants in conserved motifs of the CC and NB-ARC domains had no effect on the cell death phenotype compared to wild-type, this data suggests that R3NLR does not function as a helper NLR. To explore any role for the LRR region of R3NLR, we deleted this entire domain (R3NLR^dLRR^, removing residues 549-1069). Co-expression of R3NLR^dLRR^ with R3AK and PWT3 did not show effector-dependent cell death (**Fig. 2d, S1, S8d**). This indicates that the LRR region of R3NLR is required for R3NLR/R3AK activity but does not distinguish between a role in effector perception or activation of cell death.

### R3AK acts as the executor of cell death on effector perception

Having established R3NLR does not execute cell death, we investigated R3AK function by targeted mutagenesis. R3AK has a typical MLKL domain architecture with an N-terminal 4HB and a C-terminal kinase domain. We hypothesised that the N-terminal α1 helix of R3AK may have a role in executing cell death, analogous to some helper NLRs (55). We mutated hydrophobic residues in the α1 helix (V7E and V11E, R3AK^2E^) and tested for cell death responses. We found that mutating these two residues was sufficient to abolish cell death on co-expression with R3NLR and PWT3 (**Fig. 3a, S10a**). Protein accumulation of R3AK^2E^ was similar to the wild type (**Fig. S1**). This demonstrates that the N-terminal α1 helix of R3AK is required for execution of cell death.

**Figure 3:**
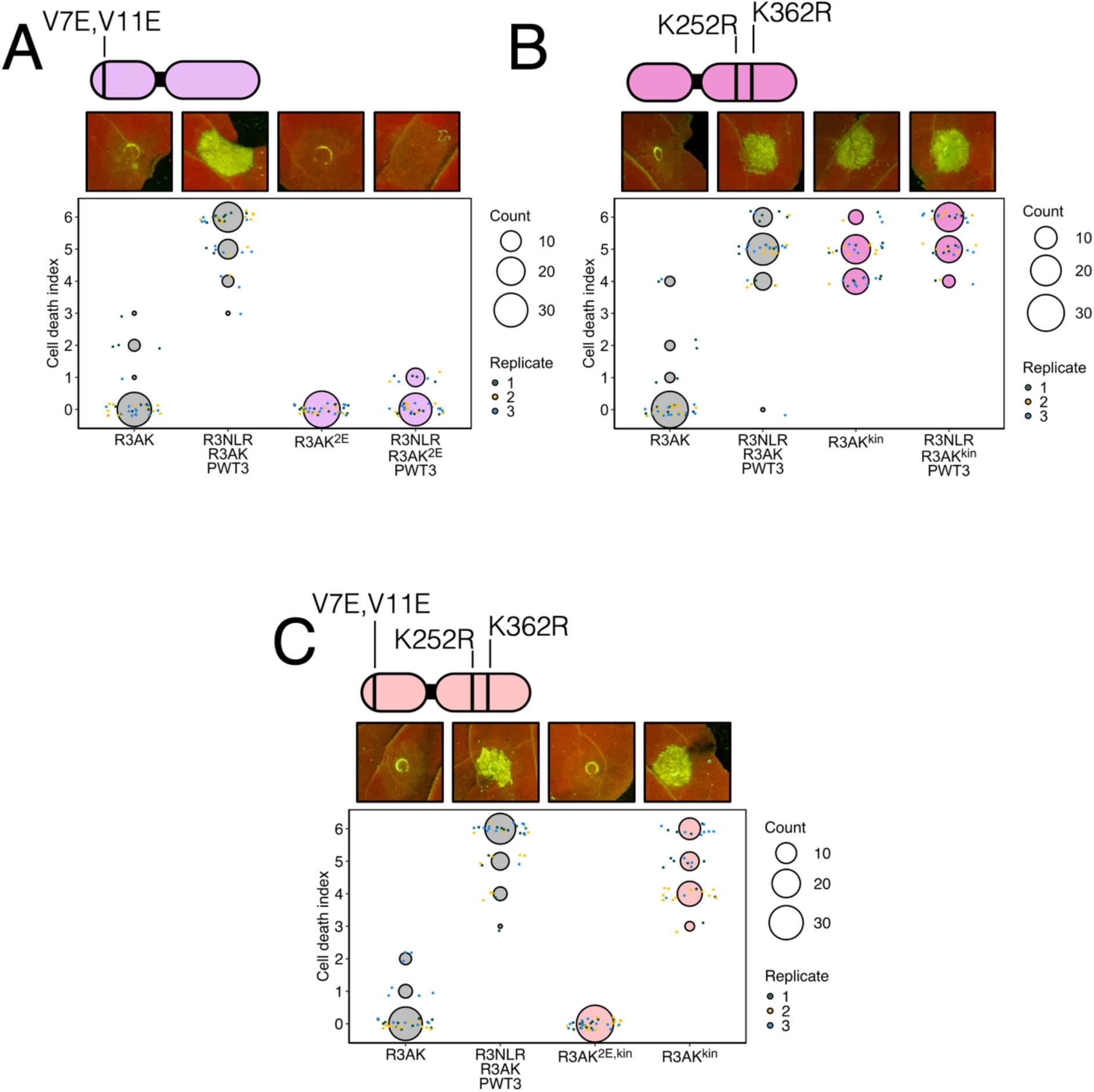
Cell death assays with mutants of R3AK show the 4HB domain is crucial for cell death signalling, and disruption to the catalytic site renders R3AK auto-active. Schematic diagrams highlighting the location of mutants within the R3AK domain structure (top) are shown alongside representative abaxial side leaf images taken under UV light (middle) and dot plots showing scoring for the cell death across repeats (bottom). (A) R3AK^2E^ N-terminal 4HB domain mutants within the α1 helix, (B) catalytic site mutant R3AK^kin^ within the kinase domain, (C) combined mutant R3AK^2E, kin^. Details of the dot plots are as for Fig. 1, with the besthr statistical analysis shown in Fig. S10.

Sequence analysis identified conserved kinase catalytic motifs in R3AK, suggesting R3AK has the potential to be a functional kinase. To investigate the role of putative ATP-binding/catalytic activity on the immune response we generated a double mutant in conserved lysine residues K252R and K362R (R3AK^kin^). Unexpectedly, when expressed in *N. benthamiana*, R3AK^kin^ caused cell death in the absence of R3NLR and PWT3 (**Fig. 3b, S1 and S10b**). To explore whether this activation required an intact N-terminal α1 helix, we tested the cell death activity of a combined mutant (R3AK^2E, kin^). Mutation of the α1 helix was sufficient to prevent the cell death of R3AK^kin^ (**Fig. 3c, S1 and S10c**). These results imply that R3AK functions as a cell death executor and may be activated through the perturbation of nucleotide binding, or through preventing its kinase activity.

### An activated R3AK forms a high-order oligomeric complex

Both MLKLs and NLRs can oligomerise in their resting state, and upon activation, form higher order complexes that result in programmed cell death (43, 46, 60–62). To probe inter-immune receptor interactions in both resting and ‘active’ states, we performed co-IP assays with epitope-tagged R3NLR and R3AK^2E^ (used to prevent cell death). Co-expression of R3NLR and R3AK with PWT3 was considered to be the ‘active’ state (as this induces cell death), while the absence of any one component would indicate a resting state (lack of cell death). We found no evidence suggesting that R3NLR interacts with itself, with R3AK or with PWT3. However, we found that R3AK oligomerises in both resting and ‘active’ states (**Fig. 4a**).

**Figure 4:**
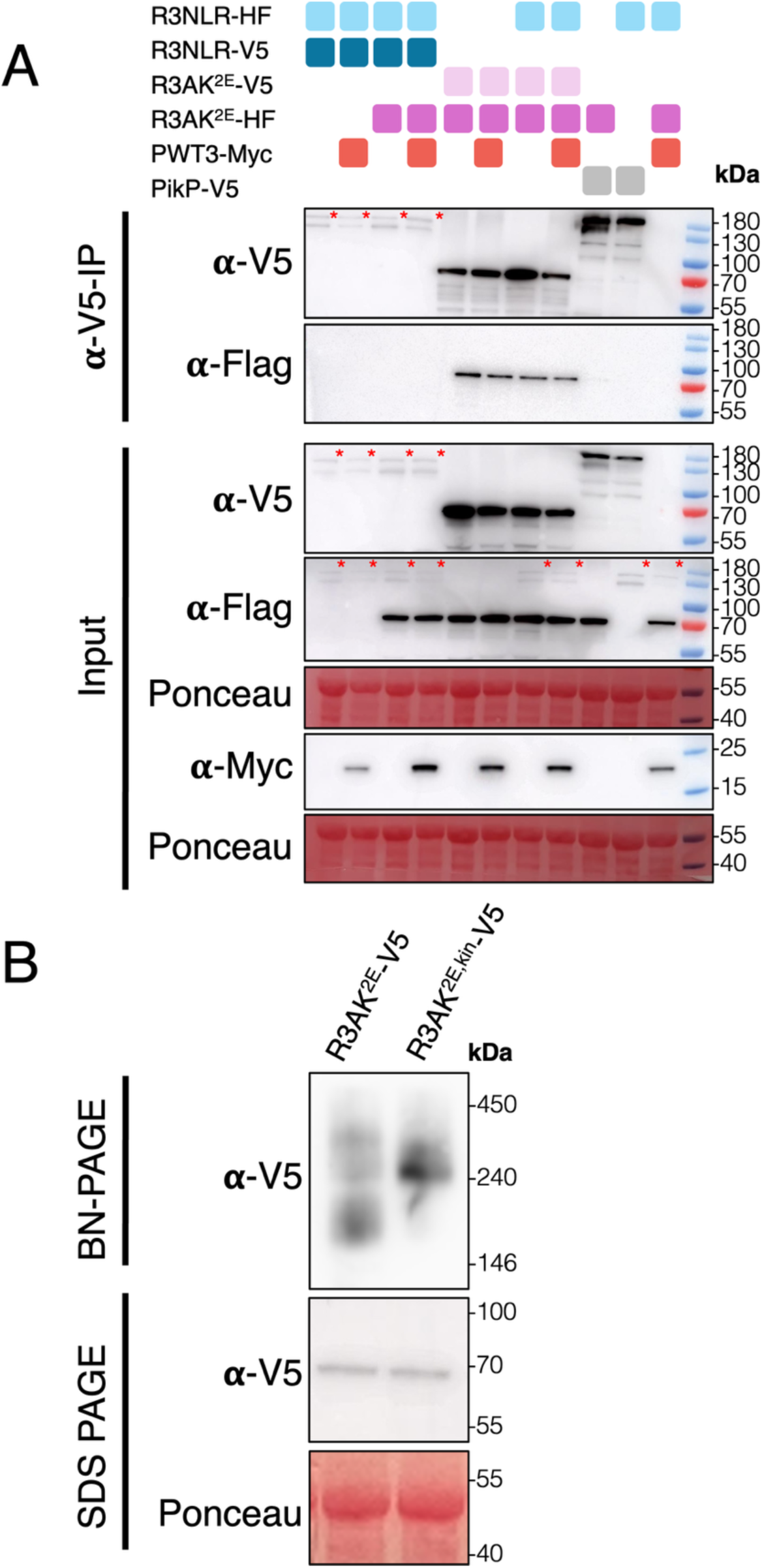
Activation of R3AK is underpinned by a change in oligomeric state. (A) Co-IP assay showing R3AK homo-oligomerisation in both ‘resting’ and ‘active’ states. All proteins were transiently expressed in *N. benthamiana* via agroinfiltration. Bottom panels (input) indicate all proteins were present before the immunoprecipitation. Anti-V5 immunoprecipitation was followed by western blot detection with relevant antibodies. Red asterix indicates the full-length band size of R3NLR. Ponceau staining was used to demonstrate even protein loading. (B) BN-PAGE assays with R3AK^2e^ and auto-active R3AK^2E,kin^ reveal an increase in size on activation. Panels show western blotting following BN-PAGE (top), SDS-PAGE (middle), both with V5 antibodies, and Ponceau staining (bottom) demonstrates equal protein loading.

By co-IP we showed that R3AK forms oligomers in both resting and active states, but it was unclear whether oligomerisation state changes on activation. To address this, we used Blue Native PAGE (BN-PAGE) assays. We observed that R3AK^2E^ (when co-expressed with R3NLR and PWT3) did not demonstrate a shift compared to the resting state, with consistent bands correlating to a molecular mass equivalent to a dimeric state observed (monomeric R3AK^2E^-V5 is ∼62 kDa) (**Fig. S11**). However, the auto-active R3AK^2E,kin^ variant demonstrated a band shift compared to the resting state, that could correspond to a tetramer (∼248 kDa) (**Fig. 4b**). Therefore, the R3AK^2E,kin^ auto-active mutant is correlated with a higher oligomeric active state able to execute cell death in the absence of other components.

Previously, AlphaFold3 (AF3) has been used to investigate the oligomeric state of helper CNLs in the presence of lipids (19, 63). As R3AK also contains a 4HB and may execute cell-death via a similar mechanism to CNLs, we used AF3 to model R3AK with 50 oleic acid molecules and scored the different oligomeric states by confidence metrics. Although overall the global prediction confidence was relatively low (highest confidence pTM = 0.55 and ipTM = 0.54), this analysis suggests tetrameric R3AK is the most confidently predicted (**Fig. S12**), correlating with results from the BN-PAGE for R3AK^2E,kin^. These predictions show strong local confidence in PAE plots for 4HB domains of R3AK to interact with each other and with lipids (**Fig. S12**). Based on these results, we suggest the active R3AK conformation is likely tetrameric, however this requires further experimental validation.

### Phylogenetic analyses reveal diverse NLR/MLKL genomic associations and multiple independent origins of monocot MLKLs

The *Pm13* gene from *Aegilops longissima* encodes an MLKL protein conferring resistance to powdery mildew (33). To investigate the phylogenetic relationship between R3AK and Pm13, we identified 360 MLKL proteins from 23 monocot genomes (15 from Poales, 2 from Zingiberales, and 6 from other monocot species) by searching for proteins that contained a kinase domain (IPR011009) fused to an N-terminal 4HB domain annotated as ‘MCAfunc domain’ (IPR045766), ‘Adapter protein Cbl, N-terminal domain superfamily’ (IPR036537), or ‘DUF1221’ (IPR010632) (**Table S4-5**). We then included Pm13 in this dataset and constructed a phylogenetic tree based on the kinase domains (**Fig. 5A**). This tree shows R3AK and Pm13 are distantly related (**Fig. 5A**). All MLKLs in clade 1 possess an N-terminal DUF1221 domain (IPR010632), whereas other MLKLs have an N-terminal ‘Adapter protein Cbl, N-terminal domain superfamily’ annotation (IPR036537), with some also annotated as ‘MCAfunc domain’ (IPR045766). None of the clade 1 MLKLs retained the examined kinase catalytic residues. We also explored whether these MLKLs have neighbouring NLR genes within 100 kbp. Some MLKLs had NLRs in this window, but there was not a clear pattern in their distribution (**Fig. 5A**). This suggests that, in addition to R3AK, other monocot MLKLs may function together with genetically linked NLRs, and that NLR/MLKL pairs may have arisen independently multiple times (**Table S3-5).**

**Figure 5:**
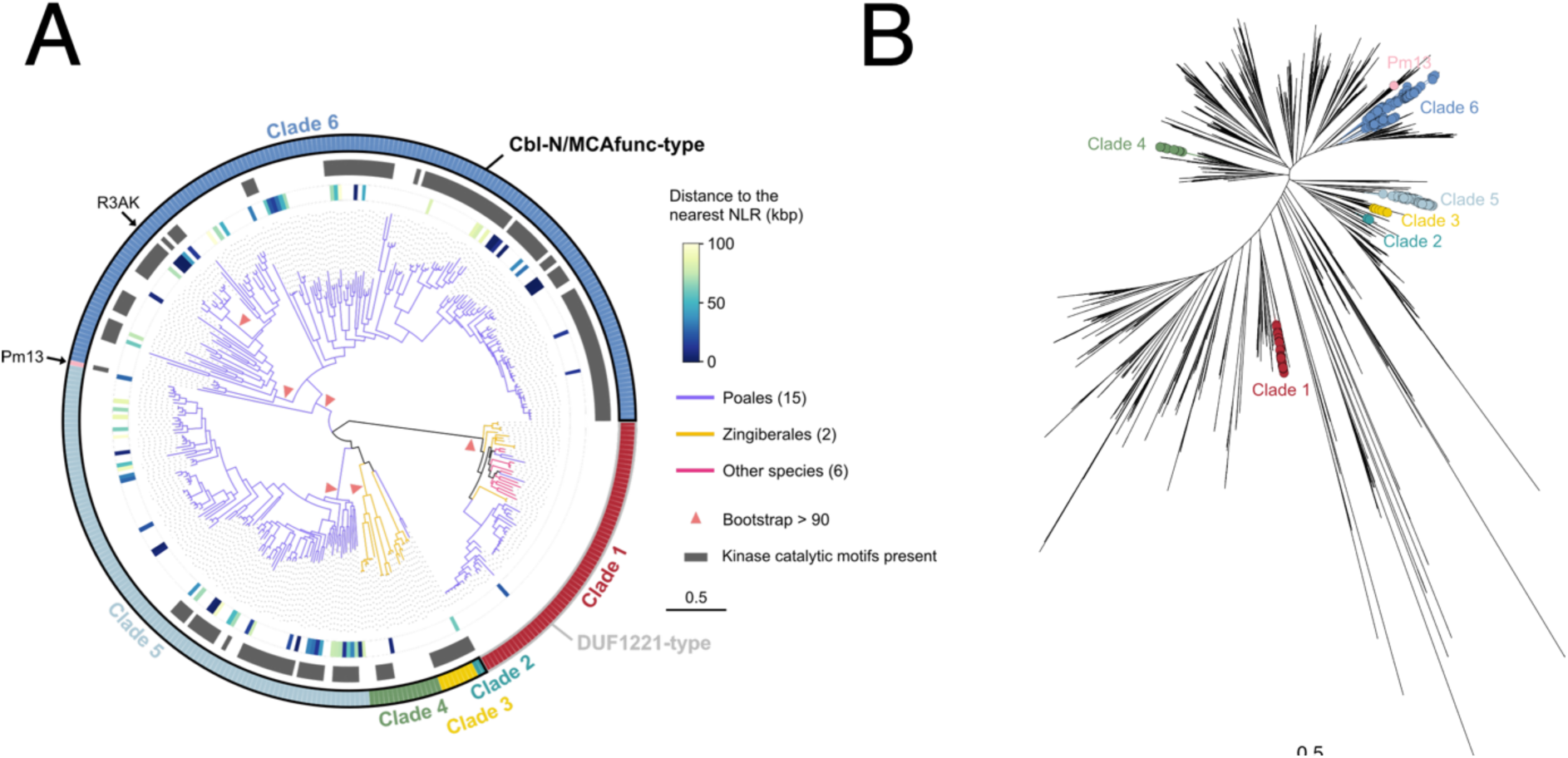
Phylogenetic analyses reveal diverse NLR/MLKL genomic associations and multiple independent origins of monocot MLKLs. (A) Phylogenetic tree of 365 kinase-domain sequences, comprising 364 domains derived from 360 MLKL proteins identified across 23 monocot genome assemblies and the kinase domain of Pm13. The conservation of kinase catalytic residues and the genomic distance, in base pairs, to the nearest NLR gene are indicated for each sequence. MLKL-associated N-terminal domain types are also indicated for each clade. The MLKL sequences were assigned to six clades, designated clades 1-6, based on their phylogenetic positions in the broader kinase-domain phylogeny shown in panel B. (B) Broader kinase-domain phylogeny used to infer the evolutionary origins of monocot MLKLs and define the six MLKL clades shown in panel A. The phylogeny comprises 4,947 kinase-domain sequences from kinase domain-containing proteins in three representative monocot species: two Poales species, *Oryza sativa* and *Setaria viridis*, and one Zingiberales species, *Musa acuminata*. The dataset also includes the monocot MLKL kinase-domain sequences shown in panel A and the kinase domain of Pm13.

To explore the evolutionary history of R3AK and other MLKLs in monocots, we extracted kinase domains from two Poales species (*Oryza sativa* and *Setaria viridis*) and one Zingiberales species (*Musa acuminata*), which was used as an outgroup to Poales. We constructed a phylogenetic tree based on the kinase domains together with the previously identified MLKL kinase domains. In this tree, the six MLKL clades (clades 1-6) are phylogenetically dispersed (**Fig. 5B**), indicating that kinase-fusion events occurred independently to form different families of MLKLs comprising different clades. The clade distribution is also shaped by taxonomic lineage. Clades 3 and 4 occur exclusively in Zingiberales (*Musa acuminata* and *Zingiber officinale*), whereas clades 2, 5 and 6 are restricted to Poales (**Fig. 5A**). MLKLs in clade 1 are the most conserved as previously identified (46) and are even retained in *Acorus*, the earliest-diverging lineage of monocots (64) (**Table S4-8**). Intriguingly, Pm13 is placed near clade 6 MLKLs but forms a separate branch (**Fig. 5**). Taken together, these results indicate that monocot MLKLs did not arise from a single ancestral MLKL but instead emerged repeatedly through independent 4HB-kinase fusion events. This pattern also suggests that R3AK and Pm13 originated from distinct fusion events.

## Discussion

Recently, a series of studies have highlighted novel diversity in disease resistance mechanisms, underpinned by KFP proteins - including acting within immune receptor pairs (27–29, 36, 38, 49, 65). In this study we investigated the mechanism of R3NLR/R3AK, a novel plant immune receptor pair, comprising an NLR and MLKL protein (49). We show that the recognition of three sequence and structurally diverse blast pathogen effectors by R3NLR/R3AK in wheat can be recapitulated in *N. benthamiana*. Mutational analysis targeting R3NLR shows the NLR does not function as a helper, but the LRR region is important for effector triggered cell death (16, 66, 67). The 4HB domain of R3AK is required for cell death and mutating two hydrophobic residues in the first α1 helix abolishes effector-dependent cell death. Moreover, activation of R3AK is correlated with a shift to a higher oligomeric state, as seen for helper NLRs. Although more complex scenarios remain possible, here we propose that R3NLR acts as a pathogen sensor and R3AK the paired helper. Our data also supports multiple kinase fusion events producing MLKLs in Zingiberales and Poales species, with some closely linked to NLRs. Overall, this work provides a new model for how NLRs and MLKLs can function together in plant immunity (**Fig. 6**).

**Figure 6:**
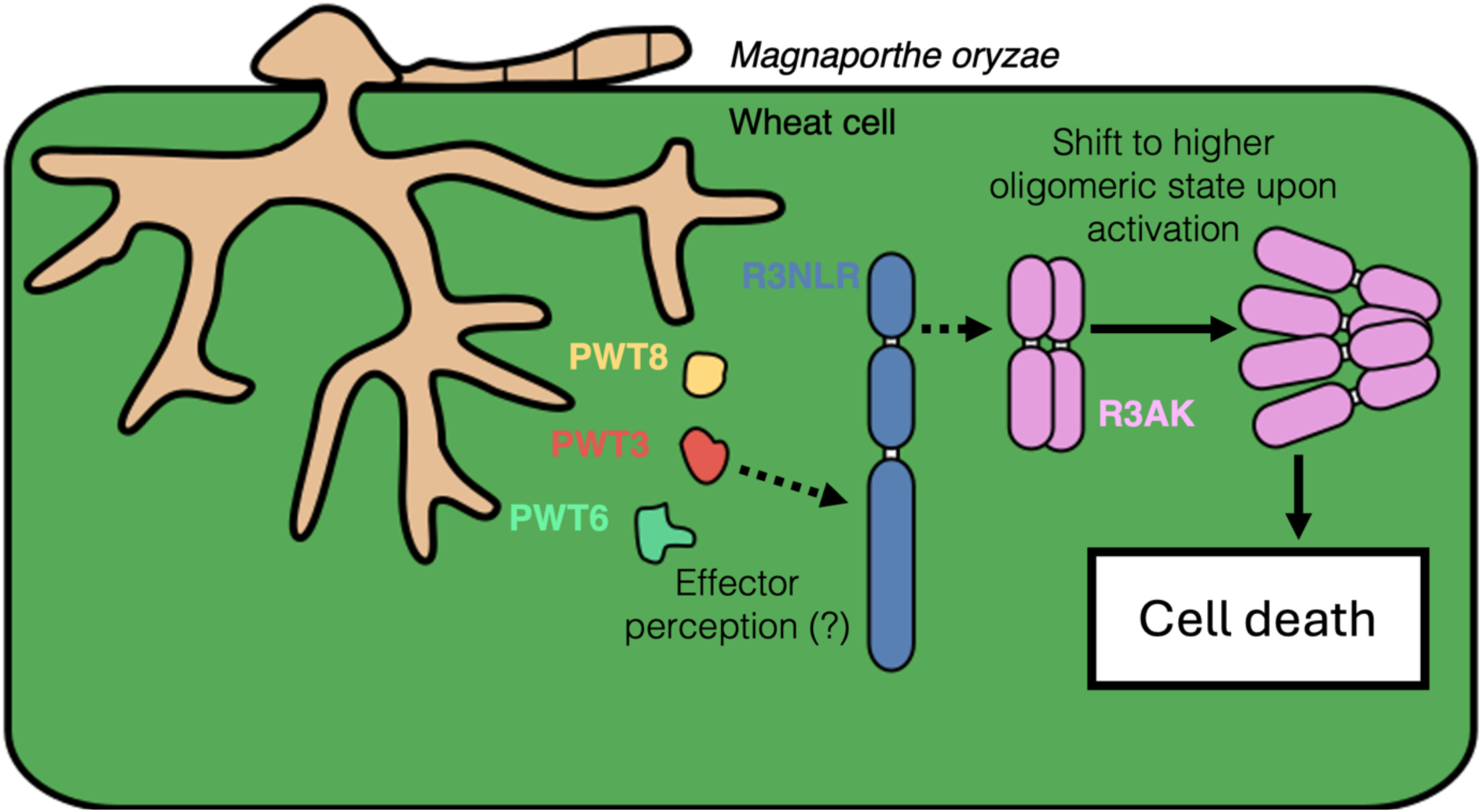
Graphical summary of the proposed R3NLR/R3AK mechanism.

Recently, the tandem kinase proteins Sr62^TK^ and RWT4 were shown to require genetically linked helper NLRs for immune signalling (29, 38). These TKPs directly bind effectors via their kinase domain, leading to the release of the autoinhibited pseudokinase domain, which in turn activates the helper NLR to cause cell death (40). The R3NLR/R3AK pair provides a novel example of a kinase fusion protein (KFP) linked to an NLR mediating immune signalling. However, unlike for Sr62^TK^ and RWT4 with their paired helper NLR (29, 38), we did not detect any association between R3NLR and R3AK in co-IP experiments. This mode of action is more similar to the Rx sensor/NRC2 helper NLR pair, where direct interactions via co-IP have also not been identified (68, 69). Nevertheless, a change in the oligomeric state of NRC2 is observed upon co-expression with auto-active sensors or through effector-dependent activation, suggesting an activation-and-release method (68–70). We observed that R3AK moves to higher molecular mass complex in the R3AK^2E,kin^ form, potentially indicative of activation, sharing similarity with Rx/NRC2. However, unlike for NRC2, where an MHD mutant of Rx is sufficient to activate signalling (71, 72), co-expression of the R3NLR^AAA^ mutant with R3AK was insufficient to alter the cell death phenotype.

Mutation of key residues within the ATP-binding site of the R3AK kinase domain results in auto-active cell death. This is similar to some NLRs, such as the flax M protein, where mutations at the ATP binding site favours an active state (57, 73). Our co-IP and BN-PAGE data support R3AK oligomerisation in both resting and ‘active’ states, and a shift to higher molecular mass on activation. However, a change in oligomeric state was not observed with effector triggered activation, only with R3AK^2E,kin^, perhaps due to reduced stability. This has previously been observed for the NLR, MLA13, where a higher oligomeric state was only observed for MHD mutants, and not effector triggered activation (74). Interestingly, ATP-binding mutants of human MLKLs (hMLKLs) favour different oligomeric states (44, 75). Resting state hMLKLs are monomers, but mutants that perturb ATP-binding can favour a tetrameric state associated with activation (76). We also observed that auto-active cell death is prevented in the R3AK^2E,kin^ mutant suggesting that the induced oligomerisation state causes cell death by signalling through the α1 helix of R3AK, much like auto-active plant CNLs (55, 61). The above comparisons show that R3AK shares mechanistic characteristics of other established yet evolutionary distinct cell death executors, exhibiting a form of functional convergence. These examples illustrate the physiochemical constraints that may restrict possible routes for evolution of successful innate immune receptors (10, 77).

The R3AK^2E^ mutation in the α1 helix abolishes the cell death response, comparable to N-terminal ‘MADA’ mutants of NLR helpers such as NRC2 (55). The clade 1 MLKLs in *A. thaliana* function in a similar manner to CNLs through forming calcium channels after activation from upstream TIR signalling (60). Furthermore, MLKLs in mammalian systems are suggested to interact with the plasma membrane and induce membrane permeation upon activation but the exact mechanism remains unclear (75, 78). Pm13, demonstrates a distinct MLKL mechanism by executing cell death through its kinase domain, with the N-terminal 4HB (HeLo) domain thought to act as a perception domain (32). Our data suggests R3AK activation promotes cell death through perturbing membrane stability via the 4HB domain, similar to NLRs, although this requires further investigation (**Fig. 6**).

An important remaining question is how the R3NLR/R3AK pair recognises sequence and structurally diverse effectors? The LRR domain of R3NLR is required for effector triggered cell death while the CC domain is dispensable. To date, we have been unable to establish direct interactions between PWT3, PWT6 or PWT8 and R3NLR/R3AK. This may be due to interactions being only transient, or experimental conditions not being suitable. There may also be a host protein guarded by R3NLR/R3AK that is binding the effectors (12). This route to perception may explain how multiple divergent effectors can be recognised by the same receptors (79). The Arabidopsis RRS1/RPS4 NLR pair recognises sequence and structurally divergent effectors via an integrated WRKY domain (79–81), but neither R3NLR or R3AK has an integrated domain.

Our data supports that MLKLs have expanded in Poales and Zingiberales as a result of multiple independent fusion events (at least 4 events in Poales and 2 in Zingiberales)(32, 33). Our phylogenetic analysis presented here showed the 6 MLKL clades in Poales, Zingiberales, and other monocot species, indicating that the MLKL domain architecture has frequently been adapted in these plant orders. The diversity of these MLKLs (clade 2–6) contrast the heavily conserved MLKLs from clade 1 that are found throughout the genomes of most angiosperm species in low numbers (45, 46). Additionally, the NLR/MLKL pairs identified are phylogenetically scattered, an indicator of multiple independent pairing events. Nevertheless, these newly identified NLR/MLKL linkages require functional validation. This phylogenetic analysis reinforces the importance of considering linkage to non-canonical resistance genes when identifying novel disease resistance in plants.

In summary, MLKLs are emerging as a source of novel disease resistance in monocots. How established examples like the R3NLR/R3AK pair and Pm13 recognise pathogen effectors is currently unknown and requires further study, especially in the context of their potential for engineering to expand use in cereal crops. This would begin to align novel non-canonical resistance gene parings to examples of NLR engineering (82–88). The observation that R3NLR/R3AK effector-dependent cell death could be recapitulated in *N. benthamiana* also raises the potential for introducing MLKL based resistance into species other than those within the Poales and Zingiberales in the future.

## Materials and Methods

### Generation of constructs for in planta expression

For in planta expression, the N-terminal signal peptides were removed for PWT3, PWT6, PWT8 (residues 1-18 for PWT3 and PWT6 and residues 1-19 for PWT8) and AVR-PikD. All constructs were constructed using modules generated by TSLSynBio and are available on AddGene (identifier codes are indicated). Effector constructs were cloned into the pICH47742 -backbone with a AtUbi10 promoter (pICSL12015 and pICSL13005) and 35S terminator (pICH41414) and an N-terminal 4xMyc tag (pICSL30009) or C-terminal 4xMyc tag (pICSL50010) with Golden Gate cloning using *BsaI*. R3AK constructs were cloned into the pICH47742 backbone with a C-terminal epitope tag (V5 or HF tags in pISCL50012 and pISCL50001 respectively) and either a NOS promoter (pICH87633) and NOS terminator (pICH41421) or an Act2 promoter (pICH87644) and Act terminator (pICH44300) for BN-PAGE to support reduced expression for clearer results. All R3NLR constructs were generated with a E1069D mutant for Golden Gate modular cloning compatibility and were cloned into the pJK0001 backbone with a 35Sx2 promoter (pICH51288) and 35S terminator (pICH41414) C-terminally tagged with either V5 or HF tags (pISCL50012 and pISCL50001, respectively). Site directed mutagenesis was used to generate specific mutants of R3NLR and R3AK constructs and the Q5 mutagenesis (New England Biolabs) kit was used.

### In planta protein expression and *N. benthamiana* cell death assays with cell death scoring

Transient gene expression was performed by infiltrating 4/5-week-old *N. benthamiana* with *Agrobacterium tumefaciens* GV3101 (C58 (rifR) Ti pMP90 (piTiC58DT-DNA) (gentR) Nopaline (pSouptetR)) henceforth *A. tumefaciens*. *A. tumefaciens* transformed with relevant constructs were infiltrated at OD_600_ 0.1 for R3AK and R3AK mutants, 0.4 for R3NLR and R3NLR domain truncations/mutants and 0.6 for PWT3, PWT6 and PWT8, in infiltration medium (10 mM MgCl_2_, 10mM MES, pH 5.6, 150 μM acetosyringone). Additionally, the p19 construct was infiltrated in all conditions using *A. tumefaciens* at OD_600_ 0.1. In all cell death assays the total OD of agrobacterium infiltrated in each condition was normalised with *A. tumefaciens* transformed with an EV control. To confirm protein expression in planta, leaf material was collected at 3 dpi (days post infiltration) (material for R3AK western blots was collected at 2 dpi to minimise any effects of overexpression) and flash frozen in liquid nitrogen. Samples were ground to a powder before mixing with protein extraction buffer (25 mM Tris (pH 7.5), 150 mM NaCl, 1 mM EDTA, 10 % glycerol, 2 % w/v PVPP, 10 mM DTT, 1 x protease inhibitor cocktail (Sigma-Aldrich), 0.1 % Tween20). Samples were then centrifuged at 4,200 xg for 30 minutes at 4°C. SDS-PAGE and western blotting were used to identify the presence of tagged proteins using their appropriate antibody (as described below).

For cell death assays, leaves were collected 4-5 dpi and imaged from the abaxial side of the leaf under UV light. Specific images shown of cell death are representative of three independent experiments, with each experiment comprising infiltrations on 6-8 leaves from 4 separate plants. Cell death scoring was performed using the cell death index described in Maqbool *et al*., 2015 (89). Dot plots were generated using R version 4.1.2 (https://www.r-project.org) with package ggplot2 (90). The size of the larger circles represents the number of times this score was observed. All individual data points are represented as dots with independent experiments coloured differently. Statistical analysis used estimation graphics from the besthr R package (MacLean, 2019).

### Co-immunoprecipitation (co-IP) assays

Genes were transiently co-expressed in 4–5-week-old *N. benthamiana* plants using OD_600_ 0.4 for NLRs, OD_600_ 0.6 for effectors and OD_600_ 0.1 for R3AK constructs. Additionally, p19 was expressed in all conditions at OD_600_ 0.1. At 2/3 dpi two leaves were collected and flash frozen in liquid nitrogen. Frozen leaf samples were ground into powder and resuspended in cold GTEN extraction buffer (25 mM Tris-HCl (pH 7.5), 10% glycerol, 1 mM EDTA and 150 mM NaCl), supplemented with 0.1% Tween 20, 0.5% w/v PVPP, 1x protease inhibitor cocktail (Sigma-Aldrich), 10 mM DTT in a 1:2 ratio. For all co-IPs investigating the interaction between the immune receptors an altered buffer was used with higher NaCl (500 mM) to minimise non-specific binding to anti-V5-beads (Sigma-Aldrich). Extracts were clarified by centrifugation, at 4,200 xg, for 30 mins at 4°C. 70 μL of the supernatant was taken and mixed with 2x Laemmli buffer and 10 mM DTT then denatured at 70°C for 10 minutes (input samples). The remaining supernatant was mixed with anti-FLAG (Sigma-Aldrich) or anti-V5 (Sigma-Aldrich) magnetic beads and incubated for 1-3 hrs on a rotary mixer. Beads were washed 5 times with cold GTEN buffer plus 0.1% Tween 20. For protein elution from the anti-FLAG or anti-V5 beads the samples were incubated at 70°C for 10 minutes with 1:1 ratio in 2x Laemmli buffer and 10 mM DTT (IP samples). SDS-PAGE and western blotting were used to visualise tagged proteins using relevant antibodies. All co-IPs and western blots were repeated with three independent biological replicates.

### Blue Native PAGE

Proteins were transiently expressed in 4/5-week-old *N. benthamiana* plants and at 2 dpi two leaves were collected and flash frozen in liquid nitrogen. Plant material was ground into powder and resuspended with GTMN extraction buffer (10% glycerol, 50 mM Tris-HCL (pH 7.5), 5 mM MgCl_2_, and 50 mM NaCl), supplemented with 0.2% IGEPAL, 10 mM DTT and 1x protease inhibitor cocktail (Sigma-Aldrich). Samples were kept on ice and resuspended by vortexing every 2 minutes for a total of 10 minutes. Samples were then centrifuged at 13,000 xg for 15 minutes at 4°C. The resulting supernatant was treated as per the manufacturer’s instructions by adding NativePAGE 5% G-250 sample additive (Invitrogen), 4x Sample Buffer (Invitrogen) and water. Samples were then run on a Native PAGE 3-12% Bis-Tris gel (Invitrogen) with a SERVA Native Marker (SERVA). Proteins from the Native PAGE gels were transferred to polyvinylidene difluoride membranes using NuPAGE Transfer Buffer (Invitrogen) and a Trans-Blot Turbo transfer machine (Bio-Rad). Proteins were fixed to the membrane by incubating for 15 minutes with 8% acetic acid and subsequently washed with water and dried. Membranes were then re-activated with 100% ethanol to visualise the unstained protein ladder and western blots performed with relevant antibodies to detect proteins. All BN-PAGE experiments were repeated with three independent biological replicates.

### Western Blot

Proteins from a 4-20% SDS-PAGE gel with a PageRuler Prestained Protein Ladder (Thermo Fisher Scientific) were transferred to a preactivated polyvinylide difluoride membrane using a Trans-Blot Turbo transfer machine (Bio-Rad). Membranes were then incubated with blocking buffer of 5% (w/v) skimmed milk in TBS-T (50 mM Tris-HCL pH 8.0, 150 mM NaCl, 0.1% Tween-20) at room temperature for 1 hr, or at 4°C overnight while gently agitated. Membranes were then incubated with either anti-FLAG-HRP (Cohesion Bioscience, 1:3000), anti-V5-HRP (Invitrogen, 1:3000), or anti-Myc-HRP (Santa Cruz Biotechnology, 1:3000) in blocking buffer for 1 hr at room temperature. Membranes were then washed with TBS-T three times and the proteins were visualised using Clarity Western ECL substrate (Bio-Rad) in a ImageQuant LAS 500 spectrophotometer (GE Healthcare). To visualise total protein loaded, membranes were stained with Ponceau S staining solution (Thermo Fisher).

### Generation of constructs for recombinant expression in *E. coli*

To enable recombinant protein expression, the PWT8 DNA sequence, without the N-terminal signal peptide, was cloned into the pPGN-C backbone with a 3C protease cleavage N-terminal 6xHIS-GB1-3C tag (pICSL30028) via golden gate cloning using *BsaI* (91, 92).

### Protein expression and purification from *E. coli*

Expression constructs comprising 6xHIS-GB1-3C-PWT8 were transformed into *E. coli* SHuffle cells. Inoculated from an overnight culture, SHuffle cells were grown in 1L of LB media in baffled flasks at 30°C until the OD_600_ reached 0.6-0.8. Cells were then induced using IPTG and incubated overnight at 18°C. Cells were pelleted at 5000 xg for 10 mins at 4°C and resuspended in lysis buffer (20 mM HEPES (pH 8.0), 20 mM Imidazole, 5% glycerol, 300 mM NaCl, supplemented with cOmplete protein inhibitor tablets (Sigma-Aldrich) (1 tablet per 50 mL)) followed by sonication. The lysate was clarified via centrifugation at 45,000 xg for 20 mins at 4°C. PWT8 was then purified through immobilised metal (Ni^2+^) affinity chromatography IMAC using a 5 mL HisTrap FF column (Cytiva). The subsequent sample was separated using size exclusion chromatography (SEC) using a HiLoad 26/600 Superdex 75 pg column (Cytiva) and SEC buffer (20 mM HEPES (pH 7.5), 150 mM NaCL). The 6xHIS-GB1-tag was removed by incubation with 3C protease overnight at 4°C. Untagged PWT8 was separated from cleaved tags and the 6xHIS-tagged 3C protease by affinity chromatography using a 5 mL HisTrap FF column (Cytiva). This was followed by a final SEC step. Resulting sample was then concentrated by ultrafiltration using centrifugation at 4,000 xg at 4°C until the desired protein concentration was reached. Protein was then flash frozen in liquid nitrogen and stored at −80°C.

### Crystallisation, X-ray data collection, structure solution and refinement of PWT8

PWT8 was concentrated to 13 mg/ml in SEC buffer (20mM HEPES (pH 7.5), 150 mM NaCl) for crystallisation screens. 96-well sitting drop plates were prepared using an Oryx8 crystallisation robot (Douglas Instruments). Crystal plates were incubated at 20°C with crystals appearing in PEGS Suite screen overnight in 2.48 M Ammonium Sulfate and 0.1 M HEPES pH 6. To enable structure solution, PWT8 crystals were soaked in selenourea for 5 minutes prior to snap freezing. Native and derivative crystals were shipped to the Diamond Light Source (DLS, UK). X-ray diffraction data was collected at DLS beamline i04 under beamline proposal mx32728. Data reduction was performed with xia2.dials and xia2.multiplex pipelines with the scaled (unmerged) data processed with AIMLESS. The structure was solved using SAD methods as implemented in the CRANK2 pipeline of CCP4i2 (93–95). The resulting protein model was then used as a template to solve the high-resolution native dataset. The final structure was obtained following manual rebuilding, refinement and validation steps with REFMAC and COOT (96, 97). The structure was further visualised using ChimeraX (98).

### AlphaFold3 modelling

Sequences of PWT3 and PWT6 without their N-terminal signal peptides were used to predict their 3D structure using the AF3 web server (https://alphafoldserver.com/) (99). To predict the different oligomeric states of R3AK, the full-length protein sequence was predicted with 50 oleic acid molecules (to mimic insertion into a lipid membrane). Attempts were made to predict four different oligomeric states of R3AK (3, 4, 5 and 6 mers) and the data was collected from 6 independent replicates without setting a seed (19, 99).

### Phylogenetic inference of monocot MLKL proteins and kinase domains

We used a harmonized monocot genome annotation dataset generated using Helixer for 23 selected monocot genome assemblies (100). Within this dataset, the gene models of R3NLR (GCA_018294505.1_CM031180.1_013693.1) and R3AK (GCA_018294505.1_CM031180.1_000255.1) were confirmed to be correctly annotated in the *Triticum aestivum* assembly GCA_018294505.1. InterProScan domain annotations from the dataset (101) were used to identify MLKL proteins and extract protein kinase-like domain sequences for subsequent phylogenetic analyses. MLKL proteins were identified from the InterProScan results by selecting proteins that contained at least one MLKL-associated N-terminal domain and at least one protein kinase-like domain (IPR011009). The MLKL-associated N-terminal domains comprised the Adapter protein Cbl N-terminal domain superfamily (IPR036537), MCAfunc domain (IPR045766), and DUF1221 domain (IPR010632). This screening identified 380 MLKL proteins across the 23 selected genome assemblies. Downstream MLKL counts and MLKL–NLR genomic-linkage analyses were based on the retained MLKL set used for phylogenetic inference, as described below. Candidate NLR genes used to assess genomic linkage between MLKL and NLR genes were identified using NLRtracker v1.0.3 (102). For each assembly, genomic distances between MLKL and NLR genes were calculated from the GFF3 annotations. MLKL–NLR pairs were identified using maximum distance thresholds of 100 kb and 15 kb.

For the MLKL phylogeny, regions annotated with IPR011009 were extracted from the 380 MLKL proteins and combined with the MLKL kinase-domain sequence of Pm13 (33). The sequences were filtered using SeqKit v2.8.1 (103) to retain kinase-domain sequences between 150 and 400 amino acids in length. The final dataset comprised 365 sequences, including 364 kinase-domain sequences derived from 360 MLKL proteins, four of which contained two retained kinase domains, and the kinase-domain sequence of Pm13. The sequences were aligned using MAFFT v7.525 with the --localpair and --maxiterate 1000 options (104). The alignment was trimmed using ClipKIT v2.3.0 in gappy mode with a gap threshold of 0.9 (105). A maximum-likelihood phylogeny was inferred using FastTree v2.1.11 (106). The resulting tree was midpoint-rooted using Biopython v1.83 (107) and visualized using Iroki (108). The conservation of kinase catalytic residues was assessed from the kinase-domain alignment used for phylogenetic inference, using residues K252 (the catalytic lysine within the “VAIK” ATP-binding motif), H358, D360 (the Histidine and Aspartate of the conserved “HRD” motif), K362 (the conserved catalytic lysine), D378, and F379 (the Aspartate and Phenylalanine of the conserved “DFG” motif) of R3AK as reference positions (30, 109). To place the MLKL sequences within a broader phylogeny of protein kinase-like domains, proteins containing an IPR011009 domain were selected from *Oryza sativa* (GCF_001433935.1), *Setaria viridis* (GCA_005286985.2), and *Musa acuminata* (GCA_032878535.1). InterProScan annotations from the harmonized genome annotation dataset were used for sequence selection. The extracted IPR011009 regions were combined with the Pm13 kinase-domain sequence and the previously identified monocot MLKL kinase-domain sequences from the 23 selected monocot genome assemblies. The combined dataset was filtered to retain sequences between 150 and 400 amino acids in length, resulting in 4,947 kinase-domain sequences. The retained sequences were aligned using FAMSA v2.2.2 in refine mode with a UPGMA guide tree (110). The alignment was trimmed using ClipKIT v2.3.0 in gappy mode with a gap threshold of 0.7 and used to infer a maximum-likelihood phylogeny with FastTree v2.1.11. The resulting tree was visualized using Iroki (108). All commands and intermediate files are available in the GitHub repository at https://github.com/YuSugihara/MLKL_kinase_phylogenetic_analyses.

## Acknowledgments

The authors thank Phil Robinson (Scientific Photographer), Julia Mundy and Dave Lawson (Structural Biology Platform), Mark Youles and Liam Egan (TSL SynBio) for their expert help and advice in this work. The authors thank all members of the Banfield Laboratory for discussions, especially Nathan J Williams for careful reading of this manuscript. JB thanks AmirAli Toghani for valuable scientific discussions throughout and Maialen Garmendia Calvo for thoughtful suggestions to improve the manuscript.

## Author Contributions

Conceptualisation: J.W.B., Y.S., P. N., S.A. and M.J.B.; Methodology: J.W.B., Y.S., J.F.H., C.A.R. and P.P.; Software: Y.S. and P.P. Validation: J.W.B. and Y.S. Formal analysis: J.W.B, Y.S, and M.J.B.; Investigation: J.W.B., Y.S., J.F.H., C.A.R., R.Z., E.Z., I.S., P.P., P.N., S.A., M.J.B. Resources: Y.S., P.P, P.N., and M.J.B. Data curation: J.W.B. and Y.S.; Writing-original draft: J.W.B., Y.S. and M.J.B.; Writing-review and editing: J.W.B., Y.S, J.F.H., C.A.R., R.Z., I.S., P.P., P.N., S.A., M.J.B.; Visualisation: J.W.B., Y.S., C.A.R. and M.J.B.; Supervision: P.N. and M.J.B.; Project administration: M.J.B.; Funding acquisition: P.N. and M.J.B..

## Funding Sources

The authors thank the Biotechnology and Biological Sciences Research Council (UKRI-BBSRC, UK, grants BB/X010996/1, BB/V015508/1), the BBSRC Norwich Research Park Biosciences Doctoral Training Partnership (grant BB/T008717/1), the John Innes Foundation, the Gatsby Charitable Foundation, and The Biochemical Society (summer studentship) for funding.

## Data and materials availability

The PWT8 structure has been deposited in the PDB server with the code 31KJ. The monocot genome annotation dataset generated using Helixer for the 23 selected monocot genome assemblies is available from Zenodo at https://doi.org/10.5281/zenodo.20703384. All commands, scripts, and intermediate files used for the phylogenetic analyses are available in the GitHub repository at https://github.com/YuSugihara/MLKL_kinase_phylogenetic_analyses.

## Notes

### Competing Interest Statement

The authors have declared no competing interest.

