## Supplemental Figures for "A wheat immune receptor pair executes cell death through a helper MLKL"

### This PDF file includes:

Figures S1 to S12

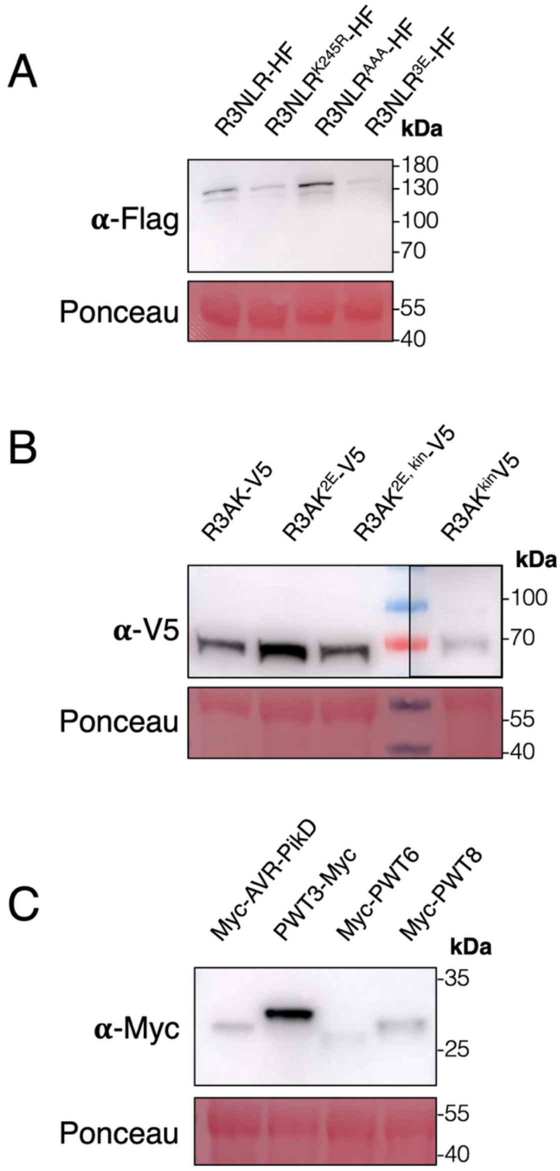


**Fig. S1: Western blot analysis to show the expression of transiently expressed proteins used in cell death assays.**

Immunoblotting after protein extraction from *N. benthamiana* tissue transiently expressing constructs used in cell death assays for (A) R3NLR proteins, (B) R3AK proteins (due to cell death, to visualise R3AK$^{kin}$-V5 the western blot needed a longer exposure time) and (C) PWT3, PWT6, PWT8 and AVR-PikD. Ponceau staining was used to demonstrate even protein loading.

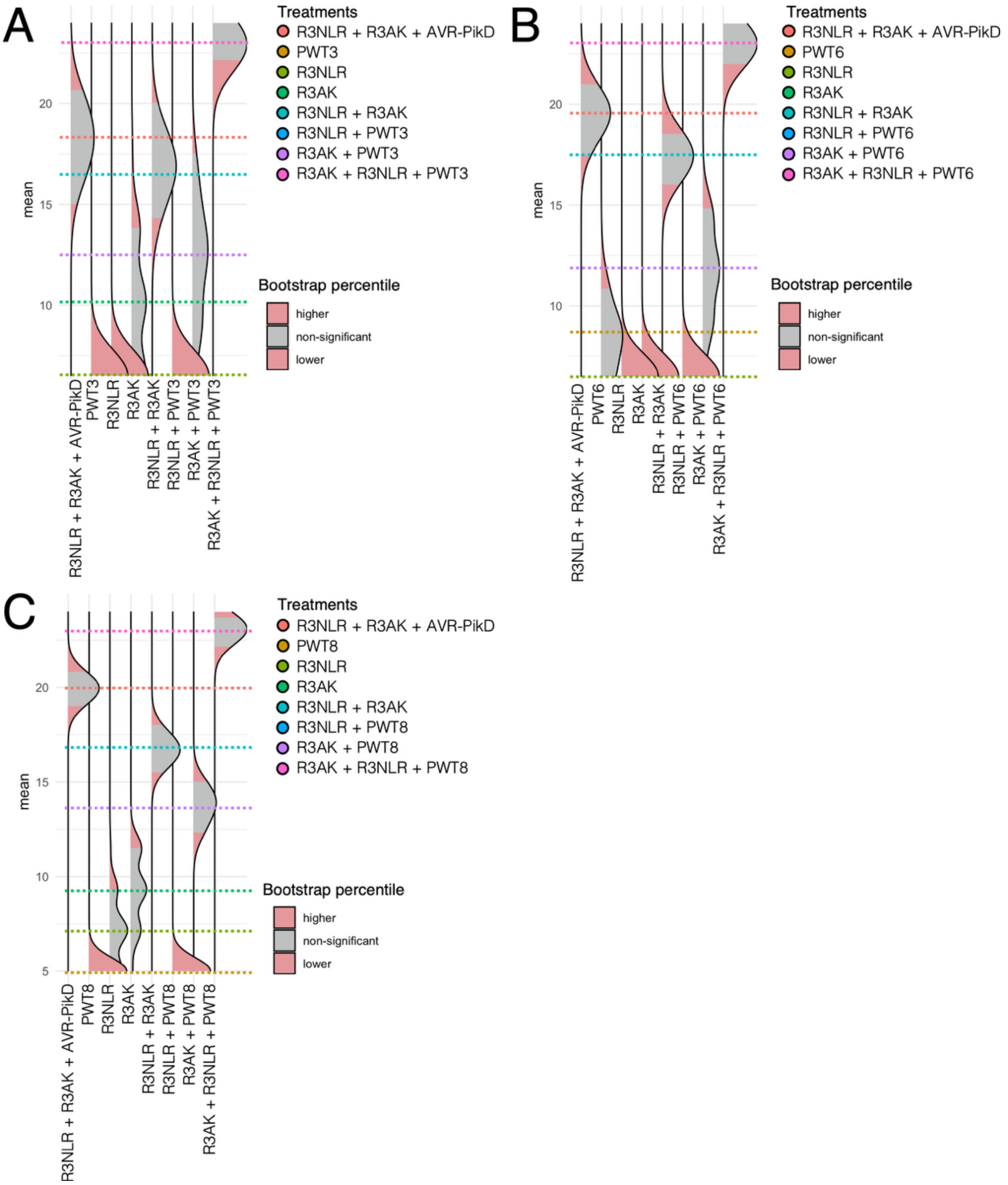


**Fig. S2: Statistical analysis of the quantification of the cell death scoring from Figure 1.** Statistical analysis using the estimation method from the 'besthr' package on R of the cell death assays shown in Fig. 1a-c, which is labelled respectively here (A-C). The distribution of 1000 bootstrap sample rank means are presented as dotted coloured lines. Red areas indicate the 0.025 and 0.975 bootstrap percentiles, whereas the grey area is the 95% confidence interval of the ranked mean. Conditions are considered significantly different if the ranked mean is outside the confidence interval of another condition (i.e. if the coloured dotted line of one condition is within or outside the red percentile of another condition).

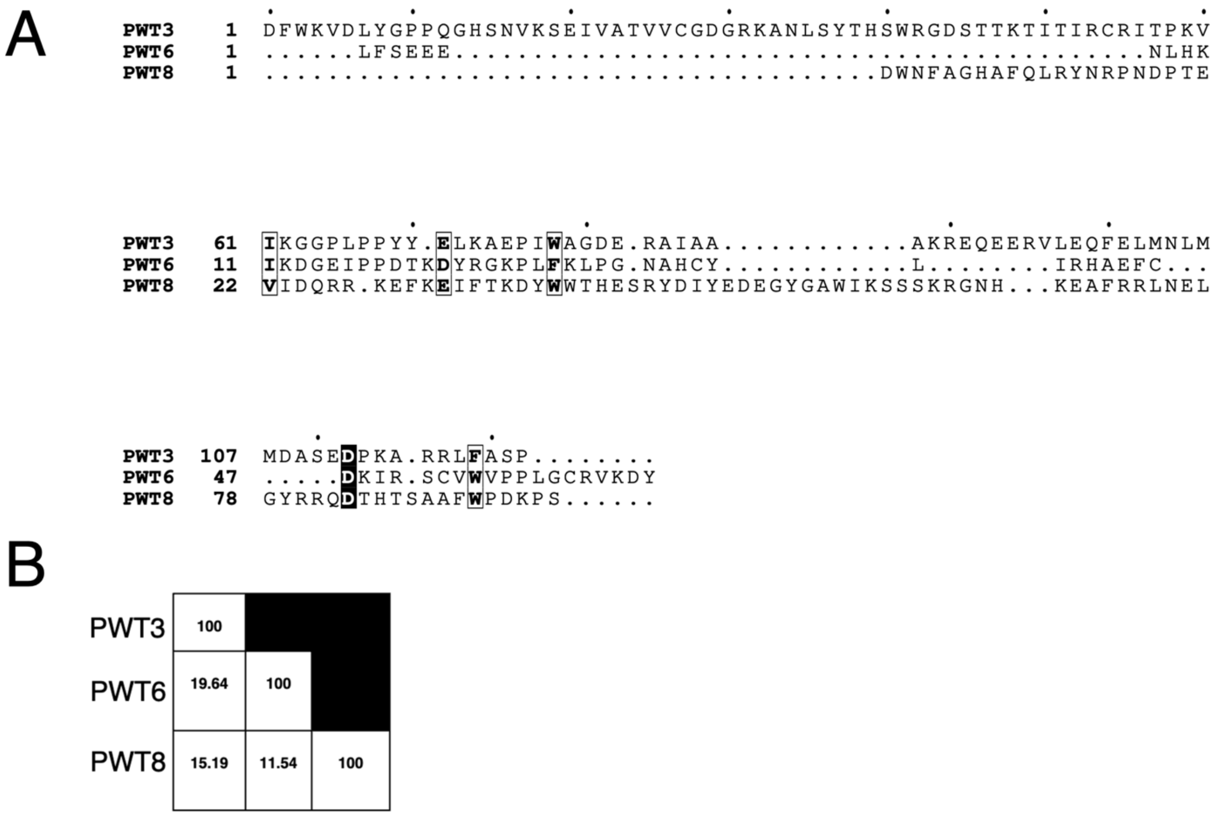


**Fig. S3: PWT3, PWT6 and PWT8 share low sequence homology.**

(A) Sequence alignment of the three effectors (with their signal peptides removed before the alignment). (B) Table indicating the sequence similarity by percentage from the alignment in A.

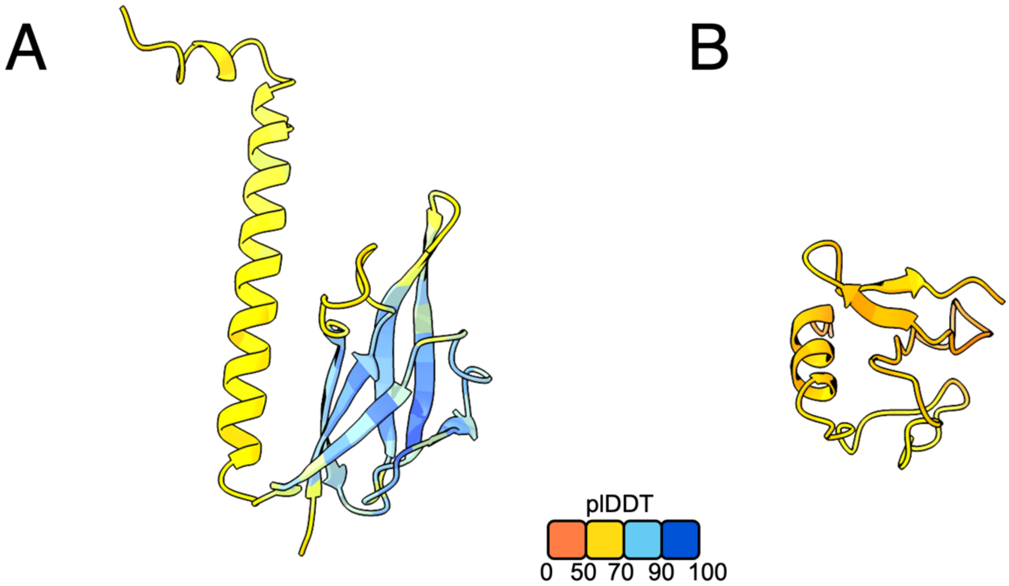


**Fig. S4: pLDDT confidence scoring of the AF3 predicted effectors PWT3 (0.56) (A) and PWT6 (0.32) (B).**

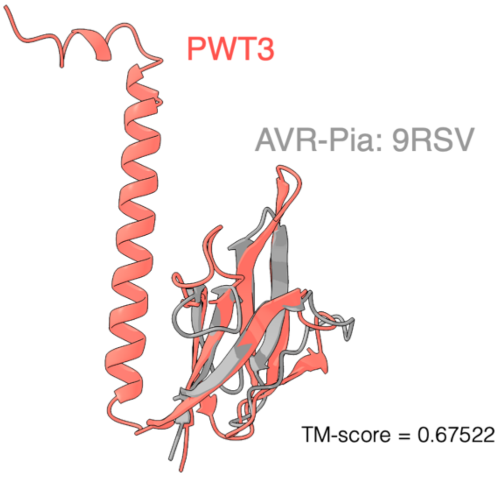


**Fig. S5: Structural comparison of PWT3 AF3 prediction to the crystal structure of AVR-Pia (PDB: 9RSV).** PWT3 is coloured in red with AVR-Pia coloured grey. The TM-score from TM-align analysis is labelled in the figure for the structural alignment.

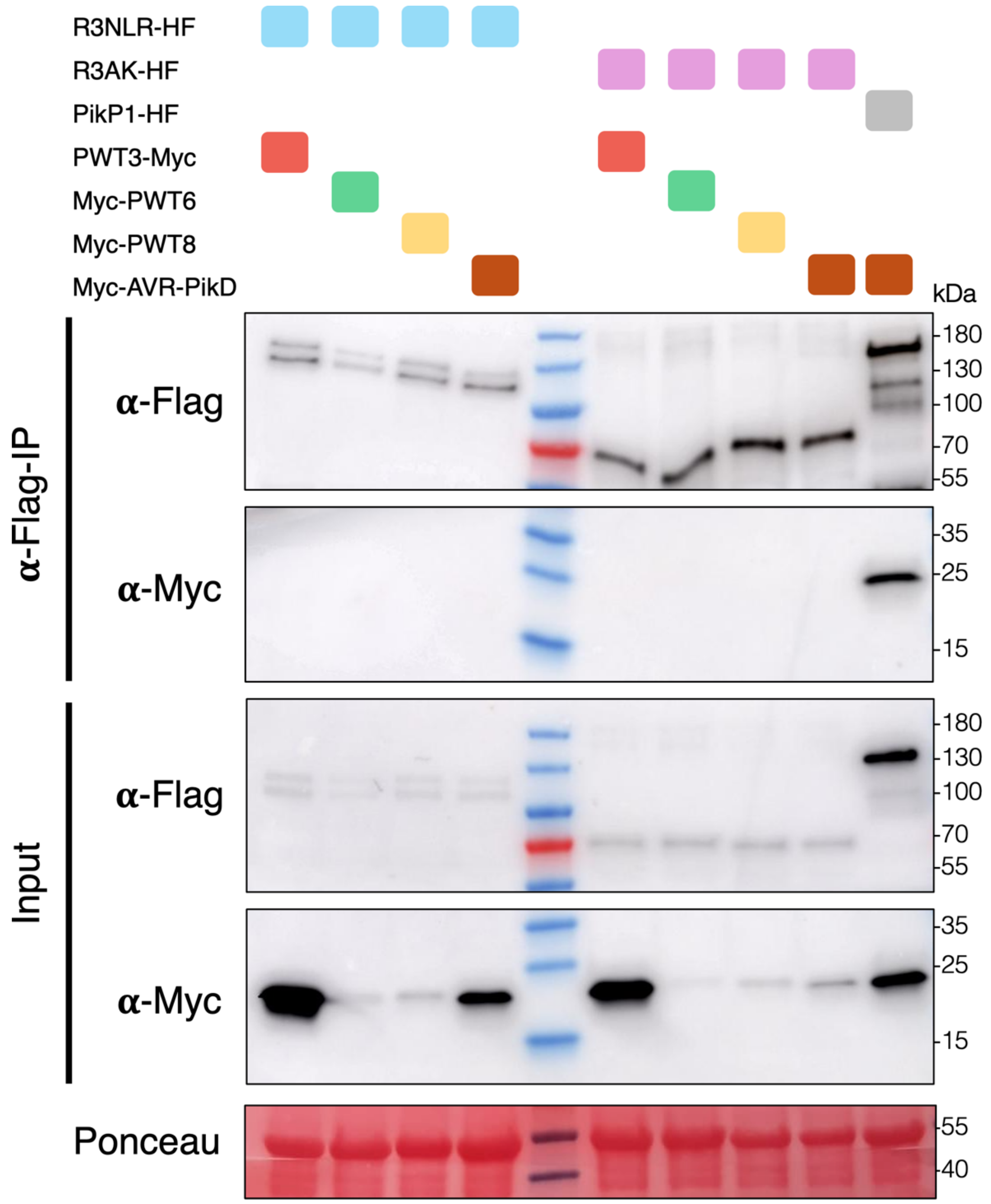


**Fig. S6: Neither R3NLR or R3AK are observed to interact with PWT3, PWT6 or PWT8 in co-IPs.**

o-IP probing for interactions of R3NLR or R3AK with PWT3, PWT6 and PWT8, shows no evidence of binding. All proteins were transiently expressed in *N. benthamiana* via agroinfiltration. Bottom panels indicate the input and that all proteins were present before the immunoprecipitation. Anti-FLAG immunoprecipitation was followed by western blot detection with relevant antibodies. Ponceau staining was used to demonstrate even protein loading.

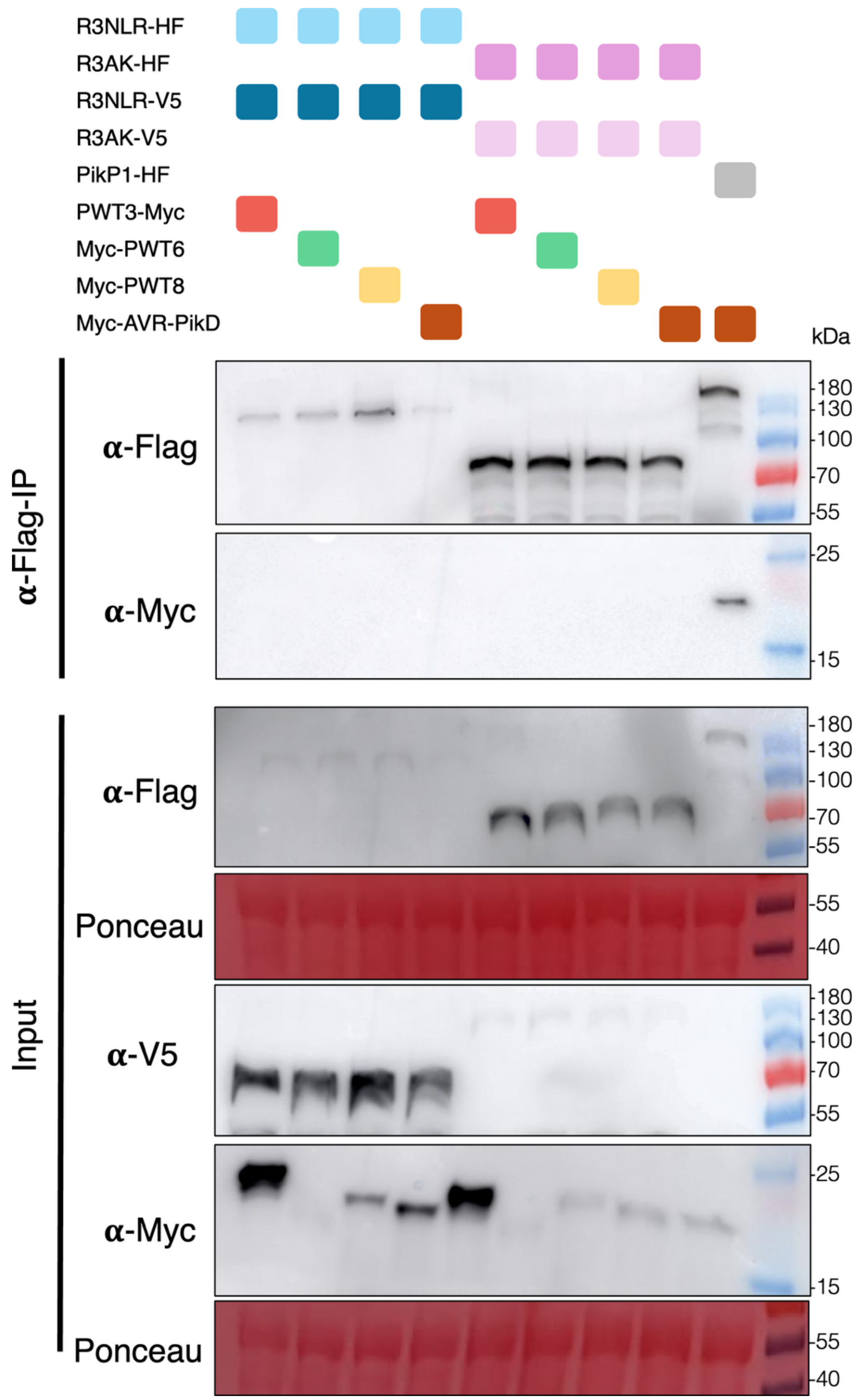


**Fig. S7: Neither R3NLR or R3AK are observed to interact with PWT3, PWT6 or PWT8 in co-IPs when co-expressed with the other immune receptor.**

Co-IP probing for interactions of R3NLR or R3AK when co-expressed with PWT3, PWT6 and PWT8, shows no evidence of binding. All proteins were transiently expressed in *N. benthamiana* via agroinfiltration. Bottom panels indicate the input and that all proteins were present before the immunoprecipitation. Anti-FLAG immunoprecipitation was followed by western blot detection with relevant antibodies. Ponceau staining was used to demonstrate even protein loading.

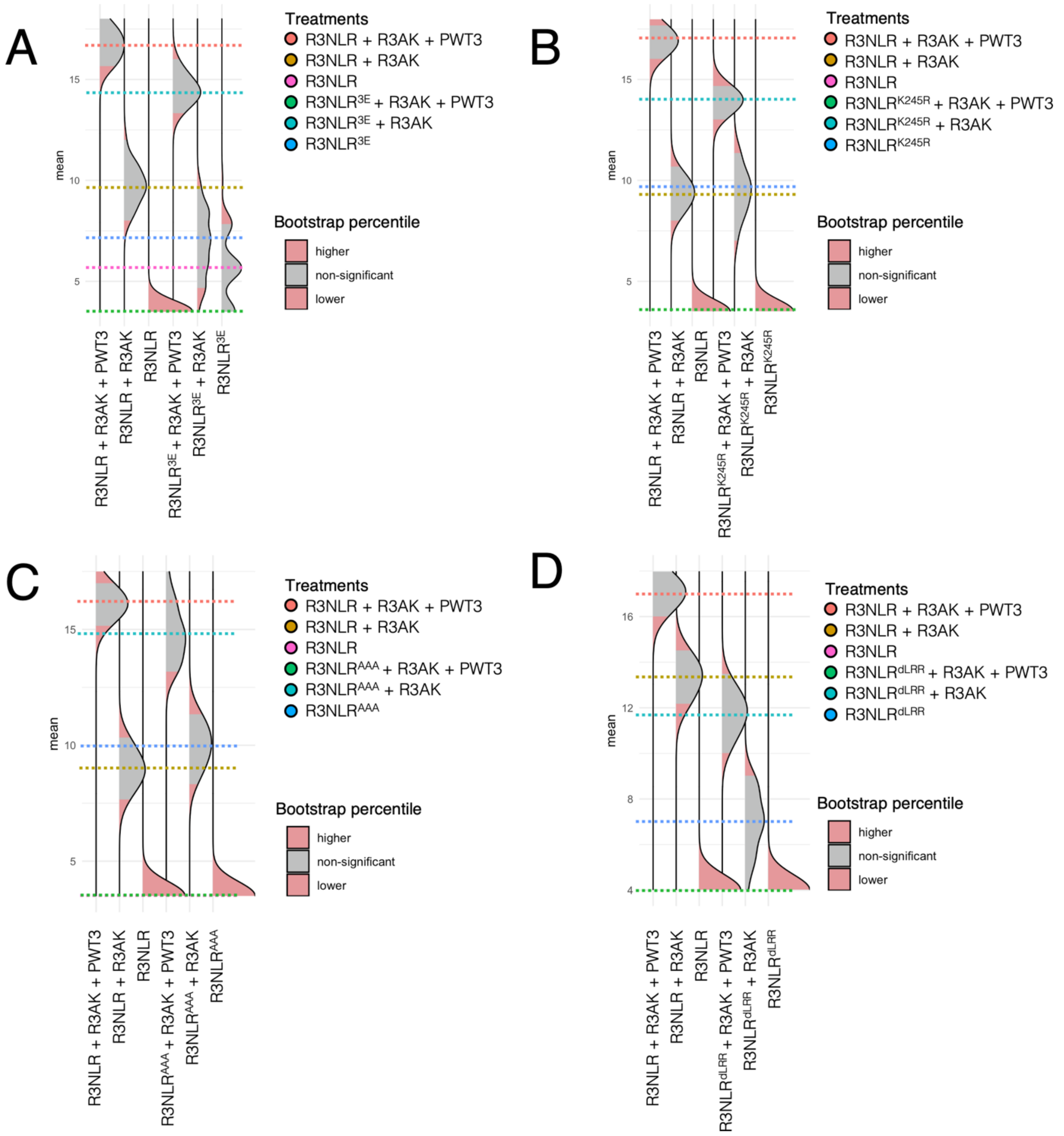


**Fig. S8: Statistical analysis of the quantification of the cell death scoring from Figure 2.**

Statistical analysis using the estimation method from the 'besthr' package on R of the cell death assays shown in Fig. 2a-d, which is labelled respectively here (A-C). The distribution of 1000 bootstrap sample rank means are presented as dotted coloured lines. Red areas indicate the 0.025 and 0.975 bootstrap percentiles, whereas the grey area is the 95% confidence interval of the ranked mean. Conditions are considered significantly different if the ranked mean is outside the confidence interval of another condition (i.e. if the coloured dotted line of one condition is within or outside the red percentile of another condition).

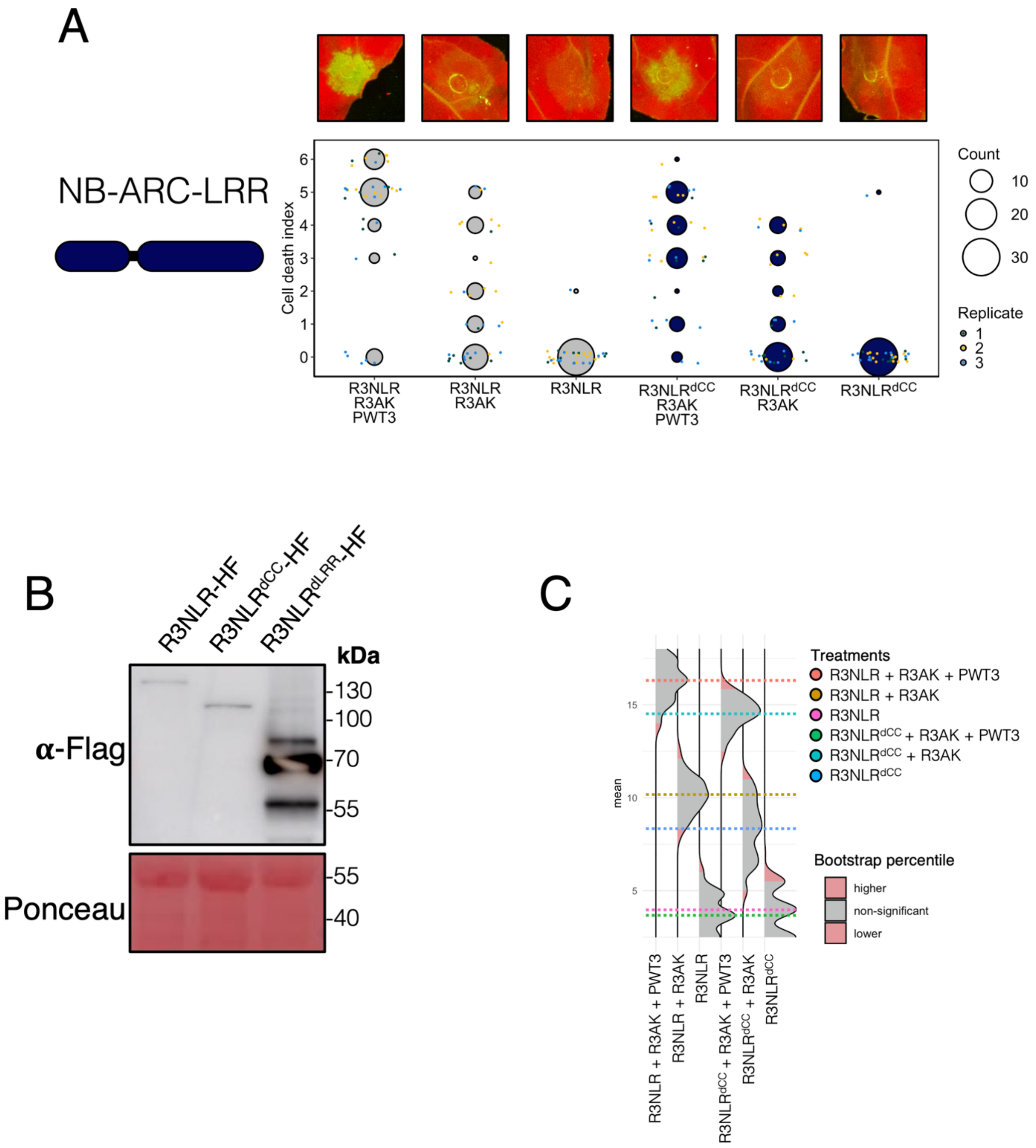


**Fig S9: R3NLR truncations show the CC domain is dispensable for cell death**

(A) Schematic diagram highlighting the domain truncation within the R3NLR domain structure (left) are shown alongside representative abaxial side leaf images taken under UV light (top) and dot plots showing scoring for the cell death across repeats (bottom). Details of the dot plots are as for Fig. 1, with the besthr statistical analysis shown in Fig. S9c. (B) Immunoblotting after protein extraction from *N. benthamiana* tissue transiently expressing constructs used in cell death assays for R3NLR$^{dCC}$ protein. Ponceau staining was used to demonstrate even protein loading. (C)

Statistical analysis using the estimation method from the ‘besthr’ package on R of the cell death assays shown in Fig. S9a. The distribution of 1000 bootstrap sample rank means are presented as dotted coloured lines. Red areas indicate the 0.025 and 0.975 bootstrap percentiles, whereas the grey area is the 95% confidence interval of the ranked mean. Conditions are considered significantly different if the ranked mean is outside the confidence interval of another condition (i.e. if the coloured dotted line of one condition is within or outside the red percentile of another condition).

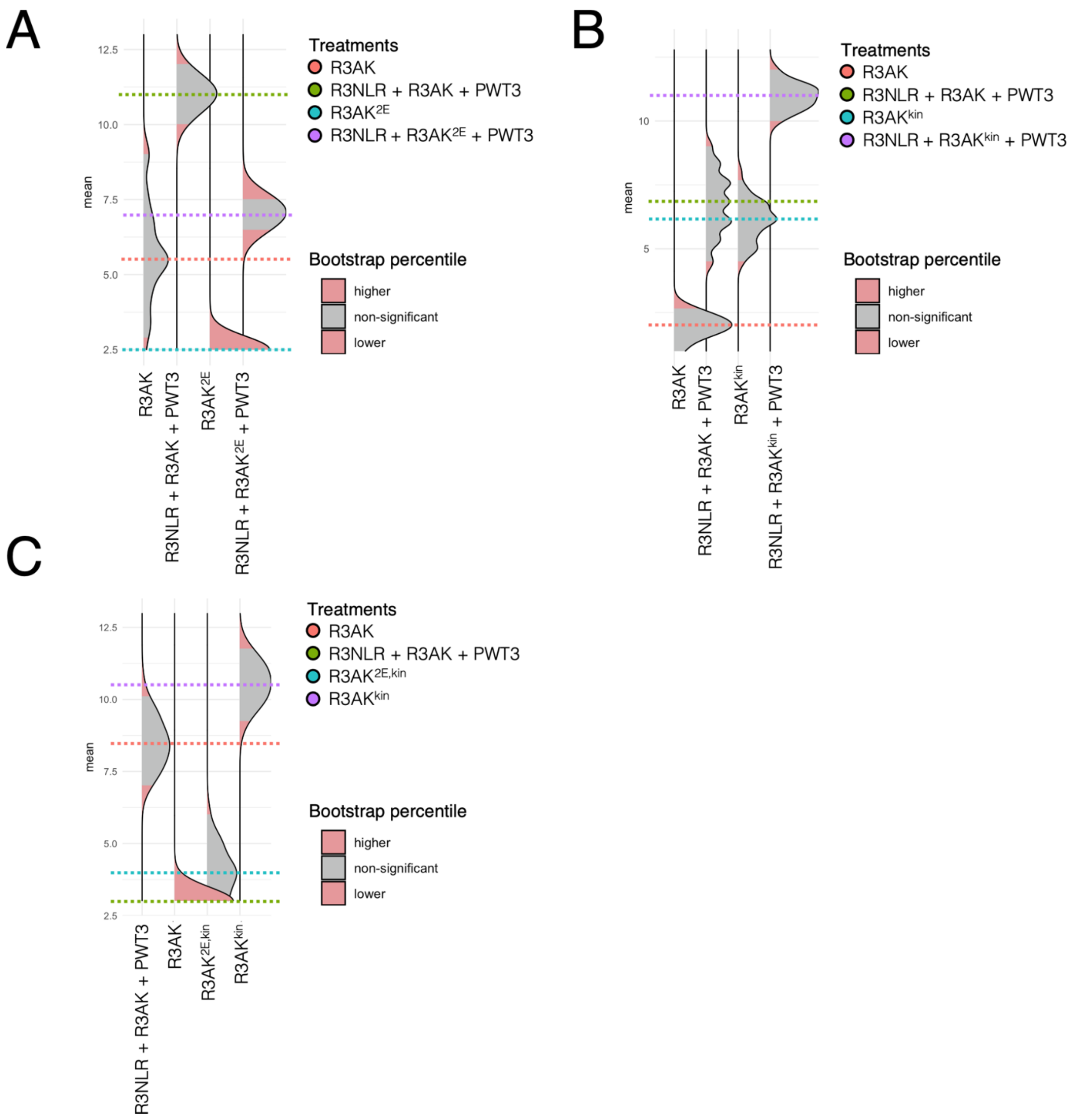


**Fig. S10: Statistical analysis of the quantification of the cell death scoring from Figure 3.** Statistical analysis using the estimation method from the 'besthr' package on R of the cell death assays shown in Fig. 3a-c, which is labelled respectively here (A-C). The distribution of 1000 bootstrap sample rank means are presented as dotted coloured lines. Red areas indicate the 0.025 and 0.975 bootstrap percentiles, whereas the grey area is the 95% confidence interval of the ranked mean. Conditions are considered significantly different if the ranked mean is outside the confidence interval of another condition (i.e. if the coloured dotted line of one condition is within or outside the red percentile of another condition).

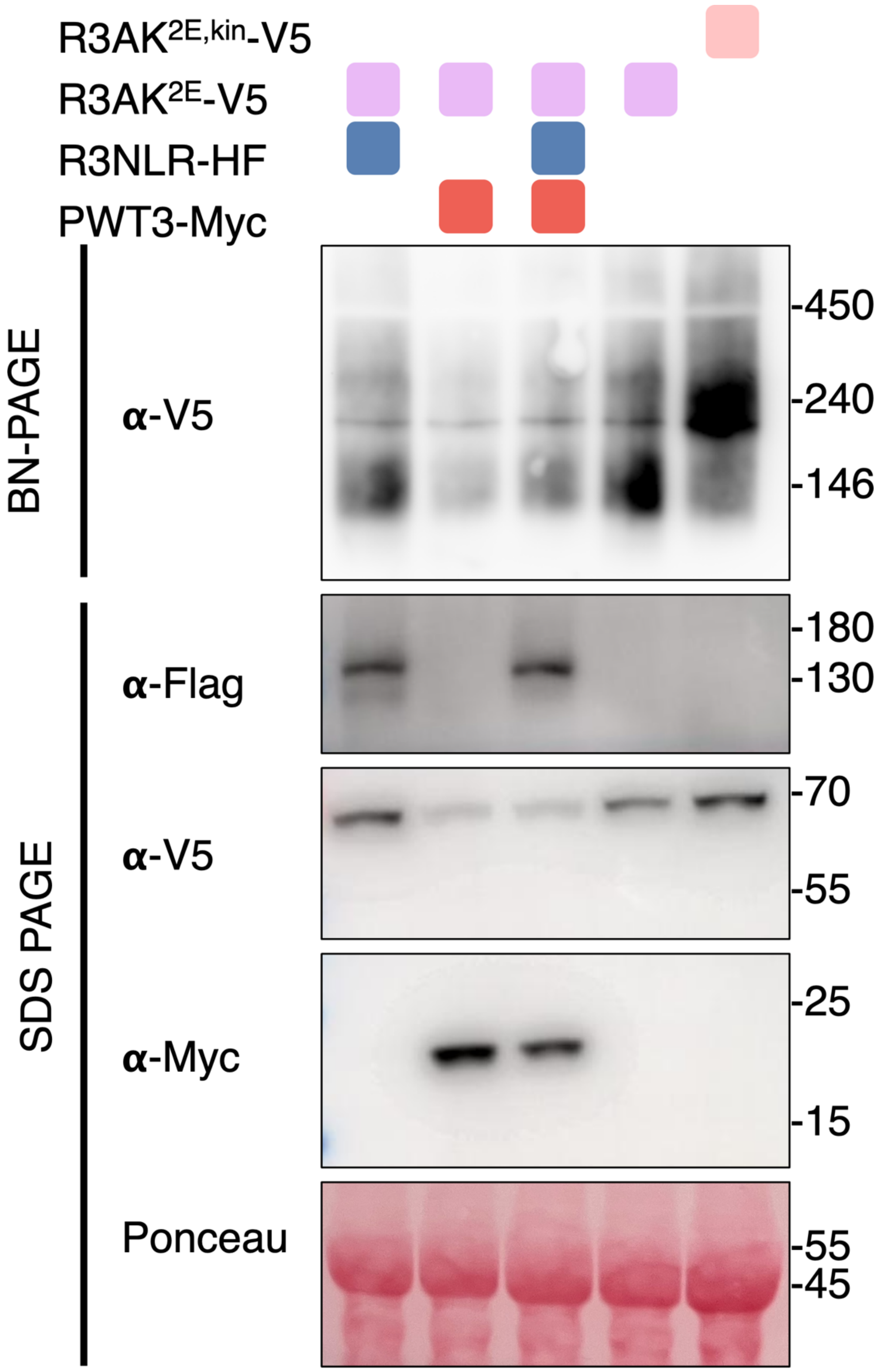


122 **Fig. S11: Auto-active R3AK exists in a higher oligomeric state than the 'inactive'**
123 **R3AK.** BN-PAGE assays with resting state R3AK$^{2E}$, 'active' R3AK$^{2E}$ and auto-active
124 R3AK$^{2E,kin}$. All proteins were transiently expressed in *N. benthamiana* via
125 agroinfiltration in three independent repeats. Top panel shows the BN-PAGE whereas
126 the bottom panels show an SDS-PAGE and western blot detection with V5 antibodies
127 and ponceau staining to demonstrate the proteins were present and expressed.

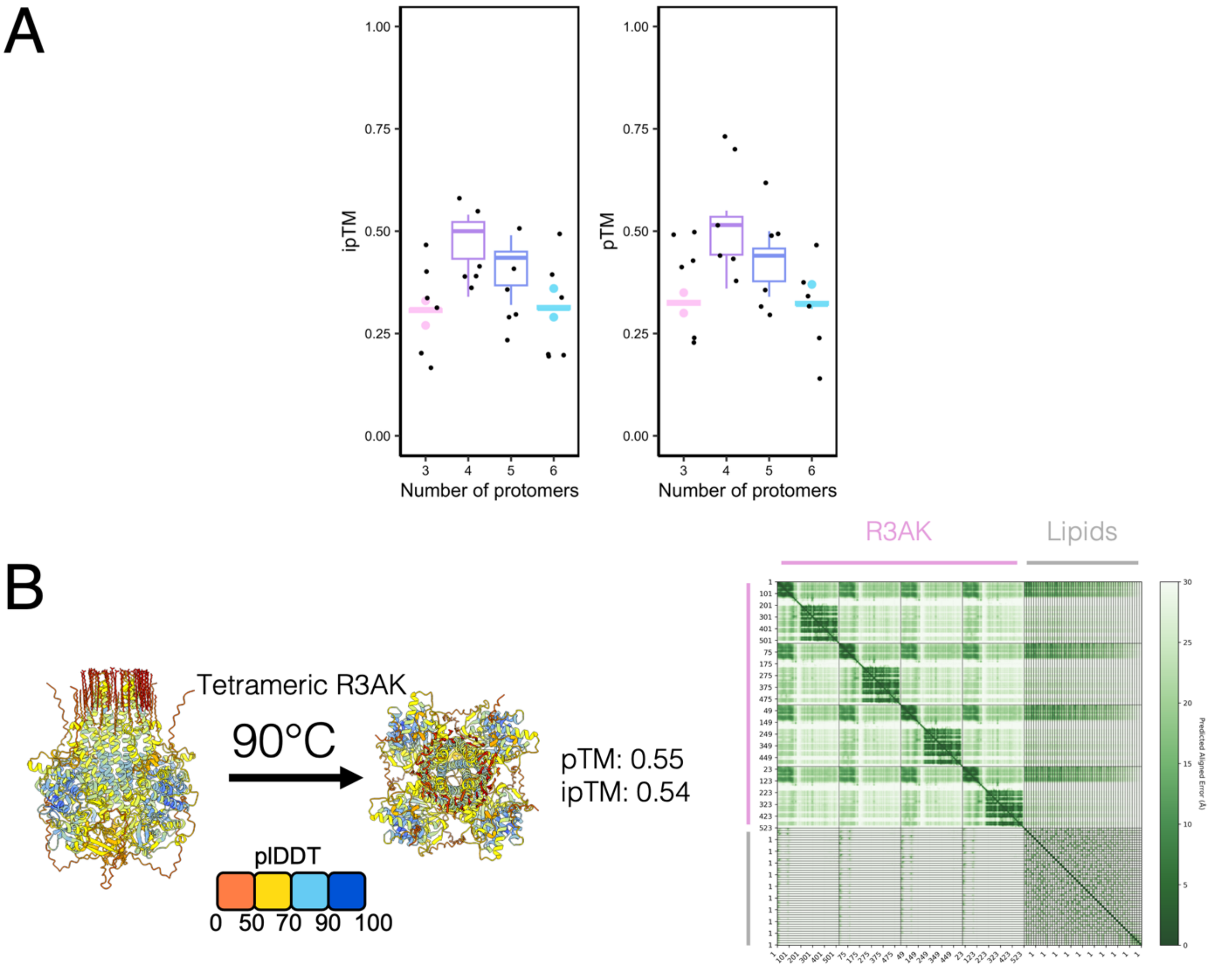


**Fig. S12: AlphaFold3 modelling most confidently predicts R3AK as a tetramer.** (A) Confidence scores from AF3 predictions with the ipTM (left) and pTM (right) plotted from 6 independent replicates using different seeds of predicted oligomeric states of R3AK in combination with 50 oleic acid molecules. (B) The structural prediction with the highest combined ipTM and pTM score from all conditions is presented with plDDT values colouring the prediction. The PAE corresponds to the prediction presented.
